# Global population structures and demographic history of *Suillus luteus*, a pine co-introduced ectomycorrhizal fungus associated with exotic forestry and invasion

**DOI:** 10.64898/2026.01.27.699563

**Authors:** Yi-Hong Ke, Anna Bazzicalupo, Joske Ruytinx, Lotus Lofgren, Thomas Bruns, Sara Branco, Brian Looney, Dai Hirose, Leho Tedersoo, Ursula Peintner, J Alejandro Rojas, Hui-Ling Liao, Jonathan Plett, Ian Anderson, Anna Lipzen, Alan Kuo, Kerrie Barry, Igor Grigoriev, Jason D. Hoeksema, Nhu H. Nguyen, Peter G. Kennedy, Rytas Vilgalys

## Abstract

Human colonization since the 19^th^ century has resulted in the global spread of pines across the Southern Hemisphere, well beyond their original northern boreal distribution. Such introductions moved not only the pines but also expanded the distribution of their symbiotic partners. Although the introduction of pines is documented through historical records, little is known about the introduction history of their ectomycorrhizal fungi, which are critical symbionts for the survival and invasion of pines. Using *Suillus luteus* as an example, population genomic analyses of 208 individuals across both native and introduced ranges showed that all introductions originated from Europe, likely mediated by human activities along with pine introductions. With the exception of North America, introduced populations were genetically differentiated from the Europe population, with varying magnitudes of population expansion in different introduced regions, often linked to forestry practices. Genetic variation within the native European population followed isolation by distance, but not in the introduced range, highlighting the disparity in the spatial genetic patterns of native versus exotic habitats. This study provides insight into the population genetics of a globally introduced ectomycorrhizal fungus whose introduction process is likely applicable to other pine-co-introduced ectomycorrhizal fungi.

## Introduction

Human activities have facilitated significant movements of plant species well beyond their natural ranges (Sakai *et al*., 2001; Meyerson & Mooney, 2007; Westphal *et al*., 2008), including the large-scale establishment of pine plantations on the foreign continents (Simberloff *et al*., 2010; Gallien *et al*., 2016). Pines (genus *Pinus* L.) are one of the most widely distributed trees across boreal forests of the Northern Hemisphere (Mirov, 1967). Since the 19^th^ century, numerous pine species have been introduced to exotic ranges in association with the global expansion of forestry and silviculture, especially in regions of the Southern Hemisphere (Simberloff *et al*., 2010; Gallien *et al*., 2016). Notable examples of such introductions include radiata pine (*P. radiata*) introduced to Australia, New Zealand, and South Africa (Wu *et al*., 2007; Burdon *et al*., 2008), lodgepole pine (*P. contorta*) introduced to South America (Simberloff *et al*., 2010; Policelli & Nuñez, 2025), and patula pine (*P. patula*) and loblolly pine (*P. taeda*) introduced to South Africa (Simberloff *et al*., 2010; Rejmánek & Richardson, 2013). The introduction of exotic pines also occurred in the Northern Hemisphere where alien Old World pine species have been planted in North America for horticultural and ornamental purposes. For example, *P. sylvestris* introduced to North America has been widely grown in Christmas tree farms, and planted along roadsides for windbreaks, shelter belts, or erosion control (Skilling, 1990).

The establishment of pine trees beyond their native ranges has not only resulted in the introduction of pine species but also expanded the distributions of their symbiotic partners (Desprez-Loustau *et al*., 2007; Pringle *et al*., 2009; Vellinga *et al*., 2009; Dickie *et al*., 2010, 2017). Pinaceae is one of the major groups of plants that form ectomycorrhizal (ECM) symbiosis with fungi (van der Heijden *et al*., 2015; Tedersoo & Brundrett, 2017; Strullu-Derrien *et al*., 2018). ECM symbiosis provides dramatic mutual benefits to fitness, such that each partner is rarely found alone under natural conditions (Selosse *et al*., 2004; Smith & Read, 2010; Policelli *et al*., 2019). The obligate nature of this symbiosis has restricted the distribution of pine-specific ECM fungi to the distribution of pines. After pine introductions to exotic habitats, the associated ECM fungal community quickly shifts to encompass introduced ECM fungi (Tedersoo *et al*., 2007; Dickie *et al*., 2010; Policelli *et al*., 2023). The ECM co-introductions can have profound consequences on local ecology, including changing the above-ground biomass, altering the balance of nutritional sequestration, increasing water use, and elevating fire risk (Chapela *et al*., 2001; Nuñez & Dickie, 2014; Dickie *et al*., 2017; Vietorisz *et al*., 2025).

Although the sources and the history of pines used in plantations are largely documented (Wu *et al*., 2007; Simberloff *et al*., 2010; Gallien *et al*., 2016), evidence of how the co-introduced ECM fungi spread is lacking. Due to their microscopic nature, the source and transportation of the ECM fungi as hyphae or spores cannot be directly observed or documented (Nuñez & Dickie, 2014; Gladieux *et al*., 2015). As a result, understanding the sources and demographic history of introduced populations of fungi has heavily relied on approaches based on molecular markers (Desprez-Loustau *et al*., 2007; Douhan *et al*., 2011; Gladieux *et al*., 2015).

Population genetics provides powerful inferences about the history and evolutionary consequences of fungal introductions. It can provide insight into the population structure of native populations, identification of introductions and their sources, subsequent admixtures arising from multiple introductions, demographic history, and signatures of adaptation (Douhan *et al*., 2011; Gladieux *et al*., 2015). For example, origins of introductions and subsequent admixed populations have been reported in several plant-pathogenic species, such as in the rice blast fungus *Magnaporthe oryzae* (Gladieux *et al*., 2018), the anther-smut fungus *Microbotryum lychnidis-dioicae* (Gladieux *et al*., 2015), powdery mildew species (Desprez-Loustau *et al*., 2018; Gur *et al*., 2021), and *Verticillium dahlia* (Milgroom *et al*., 2016). Similarly, the population genetic study on the wheat yellow rust fungus *Puccinia striiformis f. sp. tritici* has detected adaptive change toward lower capacity in sexual reproduction in introduced populations (Ali *et al*., 2010). Although these applications of population genetics have expanded our understanding of the genetic and evolutionary dynamics of fungal introductions in plant pathogenic fungi, little is known about the population genetics of introduced ECM fungi.

The slippery jack *Suillus luteus* (phylum Basidiomycota: subphylum Agaricomycotina: Class Boletales) is one of the most widely introduced ECM fungal species across the world (Vellinga *et al*., 2009; Policelli *et al*., 2019). Although native to Eurasia, *S. luteus* is reported to have been introduced to North America since 1887 (Peck, 1887), where it has expanded its host range to several North American pines such as *P. resinosa, P. strobus* and *P. contorta* (Nguyen *et al*., 2016). In the Southern Hemisphere, *S. luteus* has been broadly reported to occur under introduced pines for over 150 years in Australia, New Zealand, South America, and Africa (Vellinga *et al*., 2009; Policelli *et al*., 2019). In those locations, *S. luteus* is also associated with novel pine hosts, especially *P. radiata, P. patula, P. ponderosa*, and *P. contorta* (Policelli *et al*., 2019).

The widespread distribution and abundance of *S. luteus* in exotic pine forests make this species a valuable case study on population genomics of pine-ECM co-introduction (Hoeksema *et al*., 2020). Multiple introductions of *S. luteus* provide natural repeats to study the shared and divergent features associated with introduction within the same species. Identifying the population structure and the demographic history of introduced species is the basis for understanding further questions regarding the ecology, evolution, and management of the invasion. In this study, we explored the fundamental topics on the patterns of introductions of *S. luteus*, including (1) the overall population structures of *S. luteus* across the world, (2) the origins of the introduced *S. luteus* populations, (3) the presence of sequential introductions and admixture among geographically-close introduced populations, (4) intraspecific genetic diversity and the signatures of genetic bottlenecks in the introduced populations, and (5) the most likely process of introduction associated with observed genetic signatures. To answer these questions, we sampled *S. luteus* and its sister species, *S. brunnescens*, from the native range in Europe and Asia, and the introduced populations in Australia, New Zealand, North America, South America, and Africa. Whole genome shotgun sequencing was conducted to collect genetic data for inferring the phylogeny, population structure, and genetic diversity of *S. luteus* within and among continents to illustrate the introduction history of this important ectomycorrhizal species.

## Materials and Methods

### Specimen collection and genome sequencing

*Suillus luteus* was collected over its native range in Europe and Asia, and its introduced range in North America, South America, Africa, New Zealand, and Australia (Fig 1; Table S1). Its sister species, *S. brunnescens* (Nguyen *et al*., 2016), was also included in the sampling as the outgroup for analyses. Collections of *S. luteus* were made under native pines in Japan, Taiwan, and Central Europe. Most North American collections were associated with introduced *P. sylvestris*, though some collections from Minnesota were found under native North American Red Pine (*P. resinosa*). A complete list of collections used in this study is provided in Table S2.

**Figure 1.**
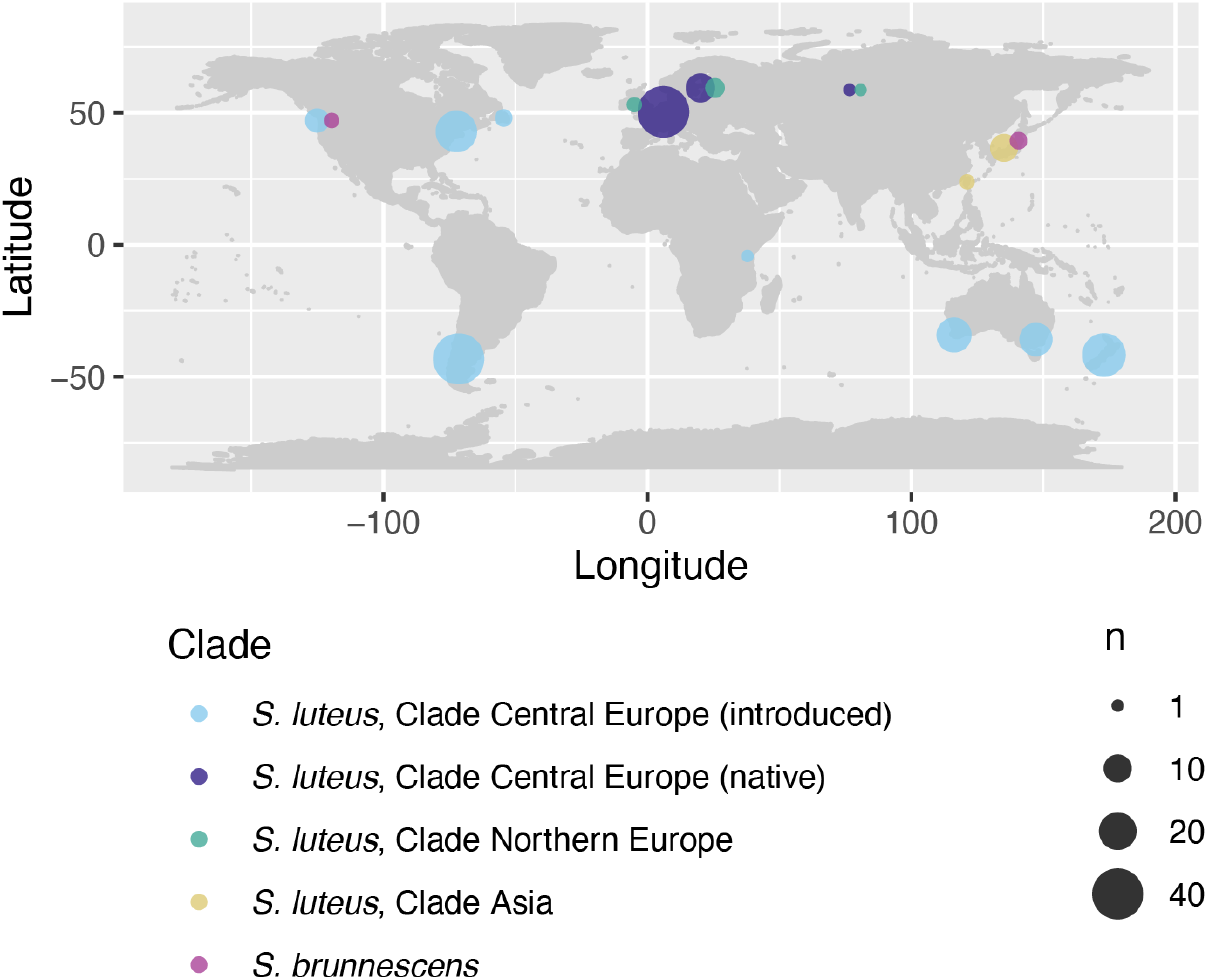
*Suillus luteus* and *Suillus brunnescens* sampling across the world. Map and sample size of populations sampled and analyzed in this study. Genome sequences were obtained from 5 *S. brunnescens* and 208 *S. luteus* collections from native and introduced ranges.

Fruiting body specimens or cultures of *S. luteus* and *S. brunnescens* were used as the sources of genomic DNA for whole genome shotgun sequencing. DNA extraction of collections from Belgium and Norway followed published methods (Bazzicalupo *et al*., 2020). For DNA extraction from tissue cultures, mycelia were grown in liquid MMN media (Marx, 1969) for 21-28 days, filtered from the media using filter paper, washed with sterile distilled water, freeze-dried, and stored at -80C until extraction. For DNA extracted from fruiting bodies, pieces of tissues cut into 5 mm cubes were preserved in 2X CTAB (2% CTAB, 1.4 M NaCl, 20 mM pH 8.0 EDTA, 100 mM pH 8.0 Tris-HCl) at room temperature until extraction. DNA extraction was conducted using OmniPrep Kit for Fungi (G-Biosciences) for both fruiting body tissues and cultured mycelia. After extraction, genomic DNA was suspended and stored in 1x TE buffer (10 mM Tris; 1 mM EDTA; pH 8.0) at -20C.

Plate-based DNA library preparation was performed for Illumina sequencing. Extracted DNA was sheared into 600 bp fragments using a Covaris LE220 focused-ultrasonicator and size selected by double-SPRI. The selected fragments were end-repaired, A-tailed, and ligated with Illumina-compatible sequencing adaptors. The libraries were sequenced in paired 2×100 or 2×150 cycles either on an Illumina HiSeq2500 or on an Illumina NovaSeq 6000. The itemized sequencing techniques and NCBI SRA accession numbers for each sample are listed in Table S3.

### Single nucleotide polymorphism discovery

Single nucleotide polymorphisms (SNPs) were determined by mapping reads to the reference genome of *S. luteus* UH-Slu-Lm8-n1 v3.0 (Kohler *et al*., 2015) and then called by *GATK* (Genome Analysis Toolkit) (van der Auwera & O’Connor, 2020). Illumina adaptors and low-quality 3’ ends under Phred score 25 in reads were trimmed using *Cutadapt* v2.1 (Martin, 2011). Reads shorter than 80 bp after trimming were discarded. Trimmed reads of each individual were mapped to the reference genome *S. luteus* using *BWA* v0.7.17 (Houtgast *et al*., 2018). To balance the depths among sequenced individuals and to save computational resources, genomes with an average depth greater than 30x were randomly subsampled to a depth equal to 30x. The likelihoods of variants of each genome were individually calculated using *GATK haplotypecaller* and then defined with *GATK GenotypeGVCFs* across all genomes. Since the reference genome contained approximately 10% annotated repeated element sequences (4,459,331 bp in 44,486,502 bp), the SNPs in the annotated repeated regions were excluded from SNPs discovery. Variants other than SNPs were filtered prior to further analyses. Hard filtering of low-quality SNPs was done following the recommendations of *GATK* documentation (QD < 2.0 || FS > 60.0 || MQ < 40.0 || MQRankSum < -12.5 || ReadPosRankSum < -8.0).

Among the 274 available genomes, we excluded genomes determined to be haploid (monokaryotic), mixed collections, mixed DNA extractions, clones of other individuals, and those cultured from spore prints which represented F1 selfed offspring (Supplementary Methods), leaving a total of 213 genomes for downstream analyses.

### Phylogeny, principal components analysis, and admixture analysis

Maximum likelihood inference of the phylogeny was conducted with concatenated SNPs using RAxML 8 (Stamatakis, 2014) with GTRCAT model without rate heterogeneity as recommended in the software manual for analyzing SNP datasets and 1000 rapid bootstrapping replicates to calculate support values of branches. *S. brevipes* Sb2 (Branco *et al*., 2015), the closest species whose Illumina-sequenced genome was available, was processed and mapped by the same procedure to the same *S. luteus* reference genomes to serve as the outgroup to root the phylogenetic tree.

To identify population structure, admixture analysis with genome-wide SNPs of *S. luteus* individuals was conducted using *Admixture* v1.3.0 (Alexander *et al*., 2009). Based on the assumption of unlinked loci in *Admixture*, SNPs were pruned if their r^2^ values to any other SNPs were greater than 0.1 in 50 SNPs windows by 10 SNPs steps using *plink* v1.9 (--indep-pairwise 50 10 0.1) (Chang *et al*., 2015) as exemplified in the program’s manual. Sites with more than 10% missing data across individuals and multiallelic sites were also removed using *plink*. Admixture analyses were run from the arbitrary numbers of populations K=2 to 8. In total 10 runs of each K value with different random seeds were conducted to ensure optimization. The runs with the highest likelihoods of each K value were considered the optimal inference of each K and used in the plots. The optimal K was determined by the 5-fold cross-validation (cv) method implemented in *Admixture*.

Principal component analysis (PCA) was used to visualize the genetic differences of individuals based on their Euclidean distances. PCA was conducted using the *dudi*.*pca* function in the R package *adegenet* (Jombart & Ahmed, 2011) using the same LD-pruned SNPs as in admixture analysis. The phylogenetic network was inferred using SplitsTree v4.18.3 (Huson & Bryant, 2006) with neighbor-net method from the same LD-pruned SNPs.

### Genetic distance and genetic diversity

Pairwise F_st_ and Reynolds’ distance were calculated to represent the genetic distance between populations, based on the same LD-pruned SNP dataset as in the admixture analysis. Hudson’s formula implemented in R package *KRIS* (https://CRAN.R-project.org/package=KRIS) was used to calculate F_st_. Reynolds’ distance (coancestry coefficient) (Reynolds *et al*., 1983) was calculated using the *adegenet* package (Jombart & Ahmed, 2011). Kinship between individuals was calculated by kinship coefficients using the KING-Robust algorithm *–kinship* in *KING* program (Manichaikul *et al*., 2010).

The significance of genetic differentiation among subpopulations in the same continent was determined by G-statistic based on allele frequency as the method described in Goudet et al. (1996) using the *gstat*.*randtest* function in package *hierfstat* (Goudet, 2005) with 10,000 randomly selected SNPs, whose significance was determined by 999 permutations. The allele frequencies of subpopulations with less than 5 individuals were not representative and discrete, so they were excluded from the test.

Isolation by distance was tested by assessing the correlation between genetic distance and geographic distance. The genetic distance matrix was calculated as Rousset’s *â* (Rousset, F, 2000; Rousset, 2008) from 10,000 randomly selected SNPs. The geographic distance was calculated from latitude and longitude coordinates with correction of WGS84 ellipsoid in *distGeo* function in R package *geosphere* (https://CRAN.R-project.org/package=geosphere). The significance of the correlation between the genetic distance matrix and the geographic distance matrix was tested using Mantel tests with 999 permutations using the *mantel*.*randtest* function in R package *ade4* (Dray & Dufour, 2007).

Genetic diversity was estimated by nucleotide diversity and segregating sites (Watterson’s θ) as described by Tajima (1989). The deviation of nucleotide diversity and the number of segregating sites from neutrality was measured by Tajima’s D (Tajima, 1989). All calculations were conducted on non-overlapping windows of 10,000 SNPs for sites having <10% missing genotypes using a customized script (scripts/tajimaD.R). The differences in genetic diversity between each introduced population and each native population were tested by Mann-Whitney U test using the function *wilcox*.*exact* in R package *exactRankTests* (https://CRAN.R-project.org/package=exactRankTests). The resultant p-values were adjusted for multiple comparisons by Holm’s method using the base R function *p*.*adjust*. The differences among introduced populations or among native populations were not included.

### Demographic modeling

Demographic modeling with *momi2* (Kamm *et al*., 2020) was used to explicitly infer the divergence history and population growth rate based on the site frequency spectrum (SFS) of genomes. *momi2* applies composite likelihood inference of models including divergence, population size, migration, and population growth rate of multiple populations at once. The mutation rate was set to 10^-7^ year^-1^ site^-1^ as used in another introduced basidiomycete fungus, *Serpula lacrymans* (Skrede *et al*., 2021). The generation time was assumed to be 10 years according to the fact that most *Suillus luteus* reproduce at trees 5-15 years old (Shaw *et al*., 2003). A simple model (A1) was applied to estimate the divergence time and population growth rate (Fig S1). The model had a single divergence event from Europe to an introduced population with a population growth rate in the introduced population. To balance the sample size among populations, the largest subpopulation in each population, namely Belgium in Europe, Bariloche in South America, and Massachusetts in North America, was treated as a representative deme for the inference. An alternative model (A1a) of the same construction except for a hard constraint of divergence time more than 1000 generations was implemented to test the possibility of long divergence from Europe to introduced populations predating human activities.

Additional models were used to test the hypotheses of sequential divergence of introduced populations. Those models contained the population in Europe and two introduced populations at once to test whether the sequential divergence scenarios better explain the divergence history than independent introductions. The first model (B1) was designed to fit the scenario of independent introductions. The model designated two introduced populations that independently diverged from the Europe population some time ago (Fig S1). The second model (B2) was designed to fit the scenario of sequential introduction. It designated that one introduced population diverged from Europe followed by the other population diverged from the first introduced population. The third model (B3) was designed to fit the scenario of independent introductions followed by subsequent admixture. It designates two introduced populations independently diverged from the Europe population, followed by a single pulse of unidirectional migration from one introduced population to the other introduced population. The population growth rates were included in both introduced populations for all models B1-B3. The detailed parameters, initial states, and optimization strategy are presented in the Supplementary Methods.

## Results

### Phylogenetic clades and global population structures

A total of 208 whole genomes of *S. luteus* and 5 whole genomes of its sister species, *S. brunnescens*, were sequenced and included in the analyses. After quality filtering, 5,741,900 SNPs were identified among the two species and the outgroup genome *S. brevipes Sb2*. When the SNP dataset was restricted to *S. luteus* individuals, the number of SNPs dropped to 2,617,891, which represented 6.54% of the non-repeated regions in the *S. luteus* reference genome.

Phylogenetic analysis of the resulting SNP dataset revealed 4 monophyletic clades consisting of *S. brunnescens* and 3 deeply divergent clades of *S. luteus* (Fig 2A). Three individuals from Japan originally identified as *S. luteus* (JS091, JS094, and JS104) were placed into a single clade with *S. brunnescens*, although these collections formed a distinct subclade from North American *S. brunnescens*. Distinct patterns of geographic distribution were observed in all three clades of *S. luteus*. The first clade consisted entirely of individuals collected from Asia (hereafter referred to as Clade Asia). The second clade included all collections from Norway (Clade Northern Europe) which was a sister group to Clade Asia. The remaining majority of European individuals originating from multiple locations in Europe formed a third clade (Clade Central Europe). Notably, Clade Northern Europe also consisted of individuals collected within the same geographic range as Clade Central Europe, such as collections made in Austria and Russia.

**Figure 2.**
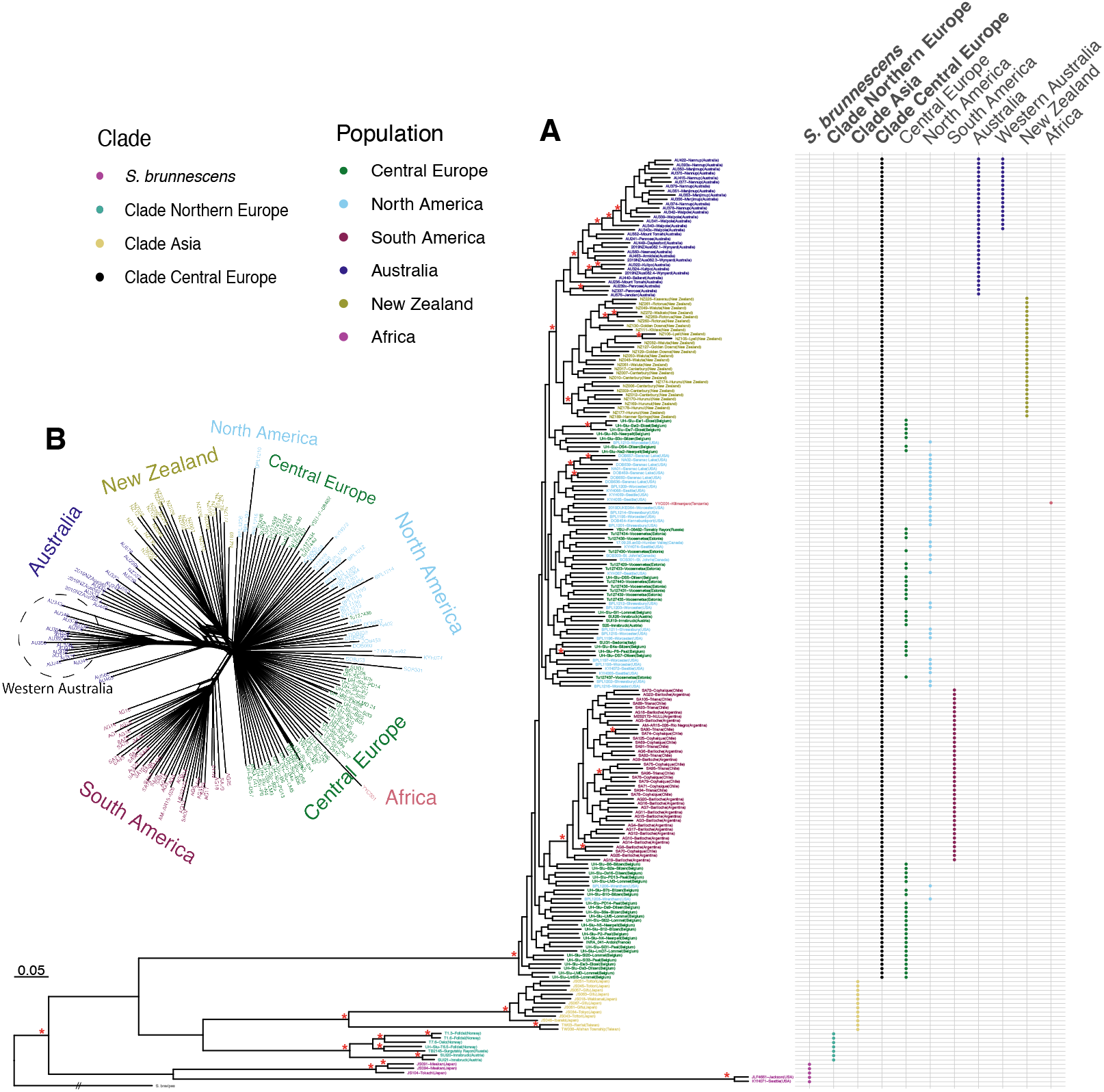
Global phylogenomic tree and phylogenetic network of the *Suillus luteus* and its closely related species. Phylogenetic relationships among the sampled individuals. (A) Maximum Likelihood phylogeny inferred from concatenated 5,741,900 SNPs of *S. luteus, S. brunnescens*, and *S. brevipes* without rate heterogeneity using RAxML. The dots on the right indicate the origins of the tip individuals. Three deeply divergent clades of *S. luteus* were present in the phylogenetic tree, namely Clade Asia, Clade Northern Europe, and Clade Central Europe. Introduced populations in Australia, New Zealand, and South America each formed a monophyletic clade diverged from Clade Central Europe. The introduced population in North America was interweaved with the native population in Central Europe. * marks indicate branches had bootstrapping support values > 95%. (B) Phylogenetic network of *S. luteus* individuals within Clade Central Europe. The phylogenetic network was inferred by neighbor-net method implemented in *SplitsTree* including both native and introduced populations. Population divergence was not observed between the Europe population and the North America population. The South America population had a clear divergence from other populations. Australia and New Zealand populations also showed divergence from other native or introduced populations, while the reticulate branches between the two populations suggest genetic exchanges were present. Western Australia (WA) formed a single cluster within other individuals in Australia as an isolated population established by founders from other parts of Australia.

The phylogenetic tree suggests that Central Europe was the main source of all the sampled introduced populations to the Southern Hemisphere and North America. In the admixture analysis, Clade Central Europe always had distinct ancestry from the other two *S. luteus* clades throughout K=2 to K=8 (Fig S2-S4; Table S4), which implied little gene flow between the other two clades and Clade Central Europe. All introduced populations had smaller genetic distances from the Central Europe population than to other clades as measured by F_*st*_ or Reynolds’ distance (Table 1). The PCA also supported that the individuals in Central Europe population and the introduced populations had a close relationship while being genetically distant from Clade Asia and Clade Northern Europe (Fig 3). Although the Asian population is geographically closer to New Zealand and Australia, it did not have observable contribution to the introduction events nor had subsequent genetic exchanges with the introduced populations. However, the possibility of introductions from multiple parts of Europe was not readily recognizable except for North America, given that the subdivision within Europe in the admixture analysis was not noticeable (K=2 to K=7) or did not share with other introduced populations (K=8).

**Table 1.** Pairwise F_st_ and Reynolds’ distance between clades and populations.

| ($F_{st}$,<br>Reynolds'<br>distance) | *S.*<br>*brunnescens* | Clade<br>Asia | Clade Northern<br>Europe | Central<br>Europe | South<br>America | North<br>America | Australia |
| --- | --- | --- | --- | --- | --- | --- | --- |
| **Clade Asia** | 0.1766,<br>0.6504 | - |  |  |  |  |  |
| **Clade Northern**<br>**Europe** | 0.1863,<br>0.6471 | 0.1374,<br>0.5967 | - |  |  |  |  |
| **Central Europe** | 0.1339,<br>0.6639 | 0.1180,<br>0.6363 | 0.1146,<br>0.6211 | - |  |  |  |
| **South America** | 0.1898,<br>0.6905 | 0.1645,<br>0.6675 | 0.1618,<br>0.6519 | 0.0246,<br>0.2447 | - |  |  |
| **North America** | 0.1471,<br>0.6653 | 0.1288,<br>0.6381 | 0.1252,<br>0.6229 | 0.0042,<br>0.1298 | 0.0268,<br>0.2524 | - |  |
| **Australia** | 0.2022,<br>0.6934 | 0.1745,<br>0.6705 | 0.1717,<br>0.6550 | 0.0261,<br>0.2507 | 0.0499,<br>0.3295 | 0.0283,<br>0.2579 | - |
| **New Zealand** | 0.1960,<br>0.6806 | 0.1695,<br>0.6560 | 0.1660,<br>0.6406 | 0.0203,<br>0.2124 | 0.0443,<br>0.3010 | 0.0220,<br>0.2199 | 0.0422,<br>0.2900 |

**Figure 3.**
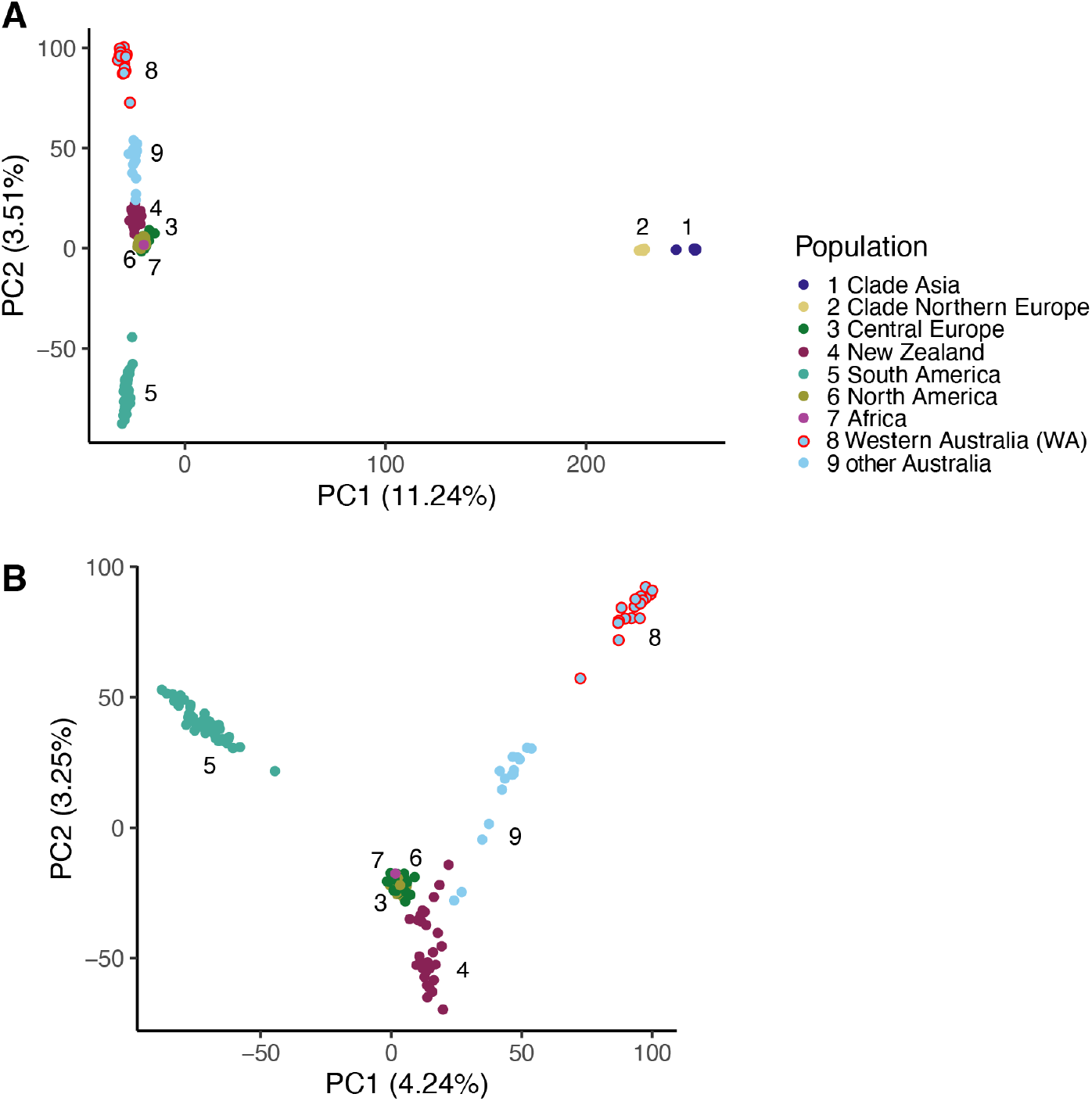
Principal component analysis (PCA) on SNPs of the sampled *Suillus luteus* individuals. The first two principal components (PC1 and PC2) based on SNPs of *S. luteus* individuals in (A) Clade Central Europe, Clade Asia, and Clade Northern Europe and (B) Clade Central Europe only. Individuals in Clade Asia and Clade Northern Europe were clearly separated from those in Clade Central Europe. Within Clade Central Europe, individuals from North America and Africa overlapped with those in the Europe population. Conversely, individuals from Australia, New Zealand, and South America did not overlap with the individuals in the Europe population. While the individuals from South America were distinct from individuals in Australia and New Zealand, the separation between Australia and New Zealand was not complete, where some individuals in Australia were closer to individuals in New Zealand than to other individuals in Australia. The individuals in Western Australia formed an isolated cluster from other parts of Australia, which positioned the individuals from other parts of Australia between the Europe population and the Western Australia population.

### Differentiation and isolation of introduced populations

Genome samples from North America showed little genetic differentiation from European populations, whose F_st_ value 0.0042 to the Central Europe population was the lowest among all introduced populations (Table 1). The phylogenetic tree and phylogenetic network also showed interleaved branches between the individuals from the two populations (Fig 2). In addition, the PCA suggested that North American individuals had close proximity to and overlap with individuals from the Central Europe population in the first two PCs (Fig 3). Lastly, admixture analysis failed to separate North America and Europe populations up to K=8 (Fig S2-S4).

Unlike North American populations, introduced populations from other regions exhibited clear population structures from the Central Europe population. Each of them formed a monophyletic clade derived from Clade Central Europe in the phylogenetic tree (Fig 2). This was also reflected by the PCA which showed genetic differentiation with no overlap between the Central Europe population and other populations (Fig 3). These results suggest that exotic introduced populations have undergone isolated evolution. Despite the high genetic similarity between Central Europe and North America, other introduced populations still had closer genetic distance to the Central Europe population than to the North America population (Table 1). In combination with no admixture and no reticulated branches in phylogenetic network with the North America population, no evidence supported secondary introduction or migration from the North America population to other continents.

In contrast, some admixture was observed between populations from New Zealand and Australia at the optimal K (Fig S2-S4), indicating shared ancestry between these populations. The phylogenetic network also showed reticulate branches between Australia and New Zealand lineages (Fig 2B). PCA further indicated few individuals from Australia appeared genetically more similar to individuals from New Zealand than with Australia (Fig 3). Such admixture suggests potential migration between New Zealand and Australia or a sequential introduction from one continent to the other, which is also reflected by their proximity in the phylogenetic tree. Other than this example between New Zealand and Australia, admixture among other continental populations was not observed.

Two independent introductions to Australia and New Zealand with subsequent migration best explain the genetic proximity of these two populations, as opposed to sequential introduction from one continent to the other. While the New Zealand and Australia populations form a monophyletic group in the phylogenetic tree (Fig 2A), F_st_ and Reynolds’ distance between the population in New Zealand and the population in Australia are still higher than either of them to Central Europe (Table 1), suggesting the populations of the two continents were independent introduction events. If these populations had resulted from a sequential introduction, the latest introduced population would have the nearest genetic distance to the intermediate population instead of the Europe population. Explicit demographic modeling on SFS also supported these as independent introductions (Fig 4C; table S5). The best model illustrated independent divergence of two populations with the Australia population receiving 37.4% of individuals migrated from the New Zealand population in the past (Model B3), which was 1,889 AIC units better than the best sequential introduction model (Model B2) and 47,266 AIC units better than the independent divergence model without migration (Model B1). This significant amount of historic admixture explains the monophyly in the phylogenetic tree and reticulation in the phylogenetic network. Furthermore, the direction of migration from New Zealand to Australia explains the presence of individuals collected in Australia genetically closer to individuals in New Zealand than other individuals in Australia in the PCA as well as the hybrid tips located in Australia in the phylogenetic network (Fig 2B).

**Figure 4.**
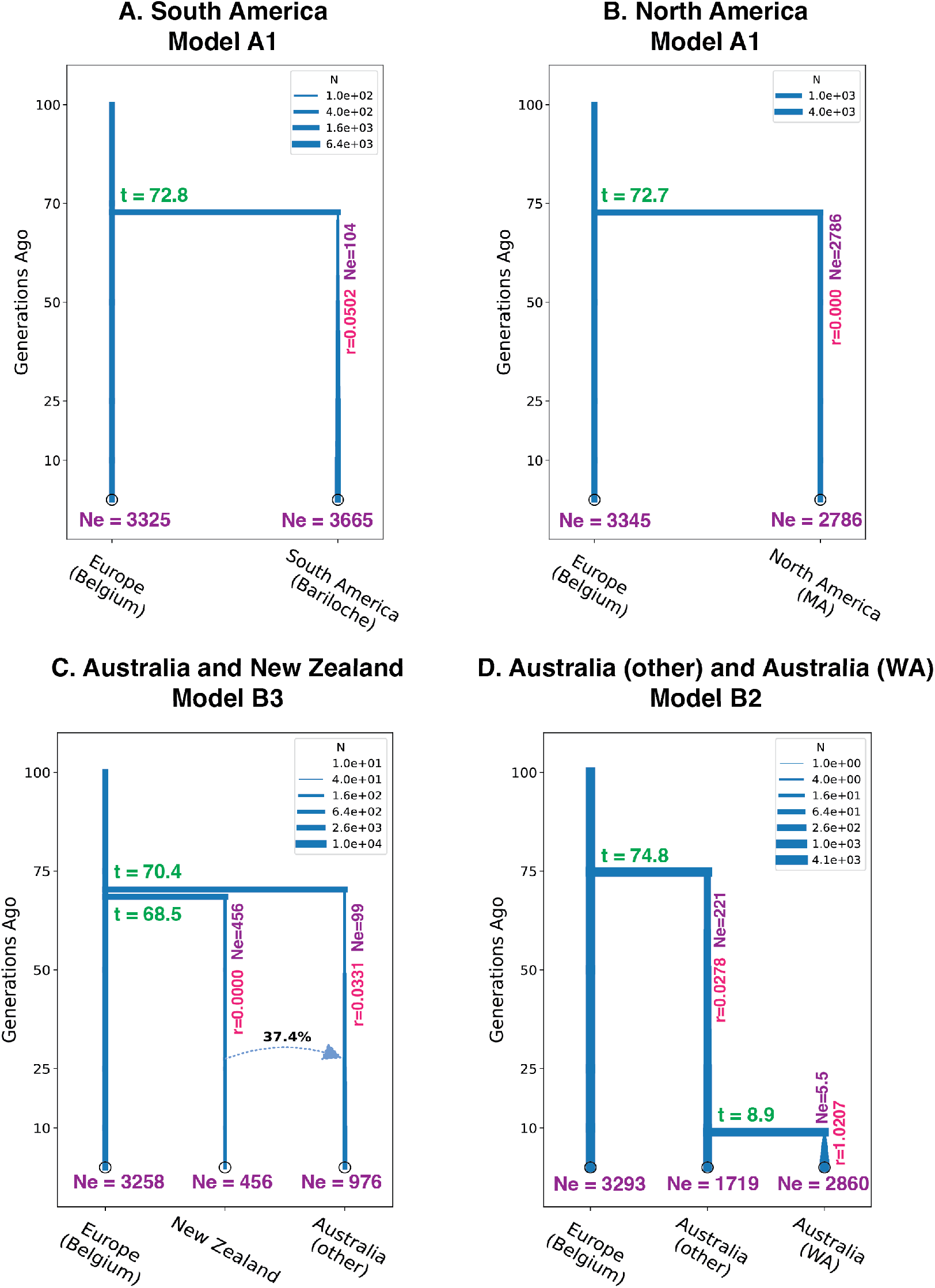
The demographic models of the most likely divergence history of each introduced population based on site frequency spectrum (SFS). The best models to illustrate the demographic history of each introduced population. **A-B** The optimized population size and growth parameters for populations of independent divergence. The inference is based on the population of Bariloche in South America and the population of Massachusetts in North America. **C-D** The optimized topologies and parameters of the divergence between New Zealand and Australia, and the divergence between Western Australia (WA) and other parts of Australia. **C** shows the better-supported scenario of independent introductions in Australia and New Zealand with subsequent migration from New Zealand to Australia (Model B3). **D** shows the better-supported sequential introduction from other parts of Australia to Western Australia (Model B2). The population size over time is presented by the width of the branches. The parameters of effective population size (Ne), divergence time (t), exponential population growth rate per generation (r), and the portion of migration in the percentage of the recipient population are labeled beside the events.

The Western Australia (WA) population is likely an isolated population from a sequential introduction from populations in other parts of Australia. The single cluster of Western Australia individuals in the phylogenetic network indicates its genetic homogeneity and supports that this population was established by a limited number of founders from other parts of Australia. The intermediate placement of individuals from other parts of Australia between the Central Europe population and the Western Australia population in PCA also hinted at the process of sequential introduction (Fig 3). Demographic modeling explicitly showed higher support for the model of sequential introduction from other parts of Australia to Western Australia (Model B2) over the independent introduction models (Model B1 and B3) (Fig 4D; Table S5).

Information on the *S. luteus* population in Africa was limited since only one collection was obtained. Nevertheless, given the fact that the individual was not nested in any other introduced population in the phylogeny (Fig 2), PCA (Fig 3), and admixture analysis (Fig S2-S4), it likely belonged to another independent introduction event unrelated to other continents, although further sampling of individuals from Africa would be necessary to confirm this hypothesis.

### Genetic diversity and effective population size

Decreases in genetic diversity were observed in all introduced populations except the North American population. The native population in Central Europe had the highest average nucleotide diversity and the highest number of segregating sites compared to the introduced populations (Fig 5). This pattern was not driven by its wider geographic range since the higher genetic diversity held even in the smaller geographic range within Belgium. Although the number of segregating sites was lower in North America in comparison to the Central Europe population, the difference in nucleotide diversity was not significant. Notably, even in the most severe loss of genetic diversity in Western Australia, the population still contained more than half of the variants as the Europe population did as measured by Watterson’s θ. Genetic bottlenecks and various magnitudes of subsequent population size expansions were observed in the introduced populations except in North America. South America, New Zealand, and Australia populations had slightly decreased nucleotide diversity and much lower numbers of segregating sites compared to the Central Europe population. This resulted in a positive Tajima’s D in all three populations and suggests a recent contraction in population sizes, likely associated with the subsampling of the parental populations and the limited numbers of founders during the introductions. Demographic modeling also supported moderate to severe genetic bottlenecks in those introduced populations. The effective population size of Europe (Belgium) was 3300 ± 50 as estimated from demographic modeling on SFS (Fig 4; Table S5). In the introduced populations other than North America, the effective population sizes of founders were smaller than that in Europe, ranging from 5.5 in Western Australia to 456 in New Zealand, which correlated well with the current genetic diversity observed in the populations. The population in Western Australia (WA) and Argentina (Bariloche), respectively, experienced dramatic 524-fold and 35-fold increases in population size since the introduction. The New Zealand population and other parts of Australia had less acute population expansions of less than 10-fold across different models. Conversely, the North America population had negligible genetic bottleneck and subsequent expansion, where the initial and current population sizes (Ne=2786) were comparable to that of Europe with zero growth rate. The estimated divergence time from Europe (Belgium) across multiple models all pointed to 75 ± 10 generations ago, which implied the similar times since introductions across the world as those of pines. The alternative hypothesis of dispersal before human activity (>1000 generations ago; Model A1a) was rejected by more than 200 AIC units worse than the optimal recent introduction models for each introduced population.

**Figure 5.**
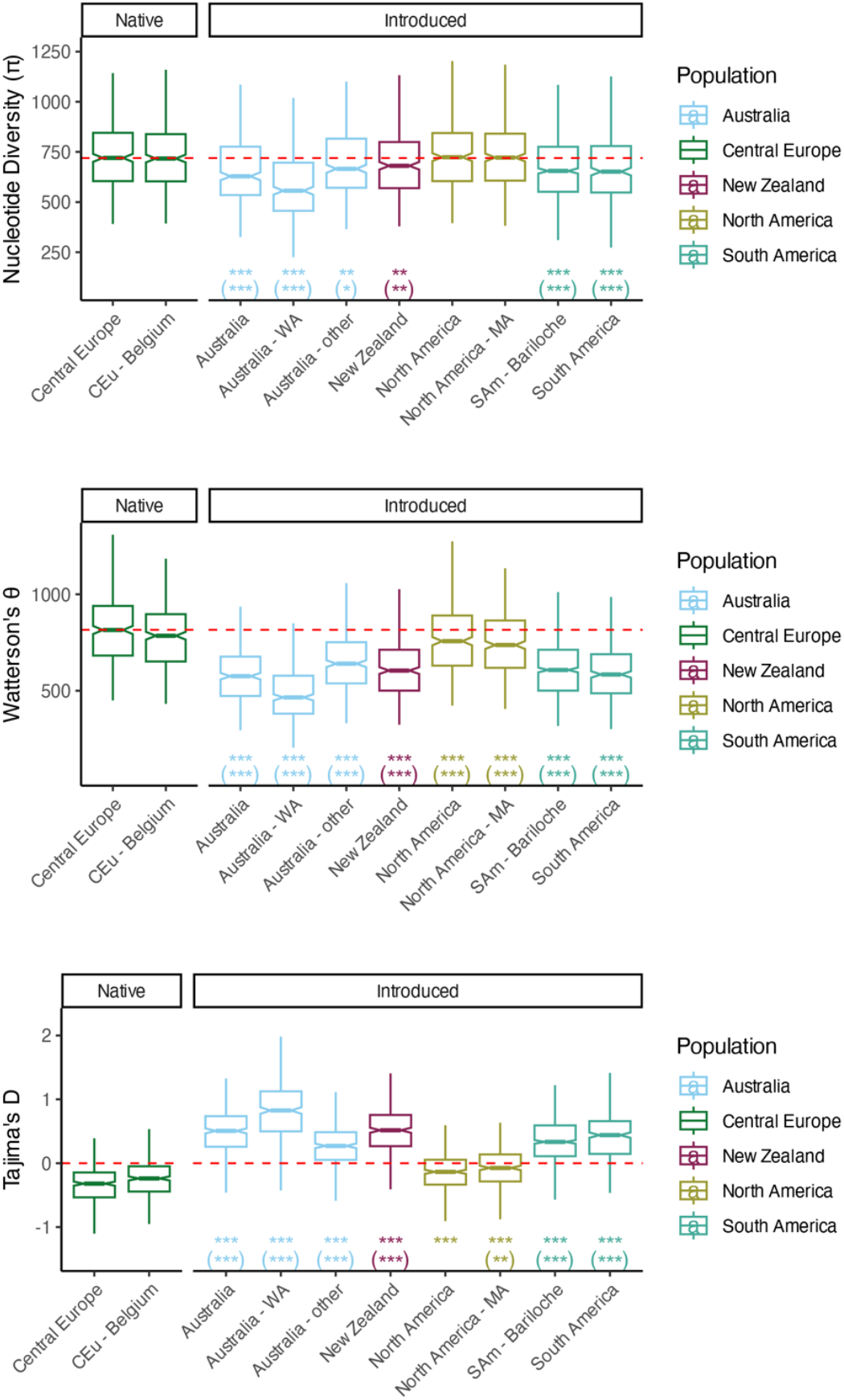
Genetic diversity and Tajima’s D of native and introduced *S. luteus* populations. Genetic diversity measured by nucleotide diversity (π) and segregating sites (Watterson’s θ) in 10,000 SNPs windows. The neutrality statistic Tajima’s D was calculated from two genetic diversity estimators. Boxplots show the distribution of the estimators in each 10,000 SNPs window. The differences between each introduced population and each native population were tested by Mann-Whitney U test whose p-values were adjusted by Holm’s method. The significance of each introduced population compared to Central Europe was labeled as ∗ for p < 0.05, ∗∗ for p < 0.01, and ∗∗∗ for p < 0.001. The comparison to the subpopulation of Central Europe in Belgium (CEu-Belgium) was shown inside parenthesis. The red dashed lines in nucleotide diversity and Watterson’s θ indicate the value of the median of the native Central Europe population. The red dashed line for Tajima’s D indicates the position of zero. All introduced populations had reduced genetic diversity, except the decrease of nucleotide diversity was not significant in North America. Tajima’s D suggests population contraction in all introduced populations except for North America. Australia – WA: Western Australia; North America – MA: Massachusetts, USA.

### Subpopulation differentiation in native and introduced ranges

The subpopulation differentiation and the spatial relationship of individuals within the same continent vary across continents. In the Central Europe population, geographic distance and genetic distance were significantly correlated (Mantel test, p<0.05; Fig 6D), suggesting a pattern of isolation by distance. Although the differentiation on PC1 of Central Europe population (Fig 6B) was driven by three individuals in Belgium which had close relationships (kinship coefficient > 0.2), the values of PC2 mirrored the geographic gradient from the west to the east of Europe, suggesting the subpopulation structure is a consequence from gradient dispersal. The same pattern persists when the remote individuals in Russia were excluded (Fig S5).

**Figure 6.**
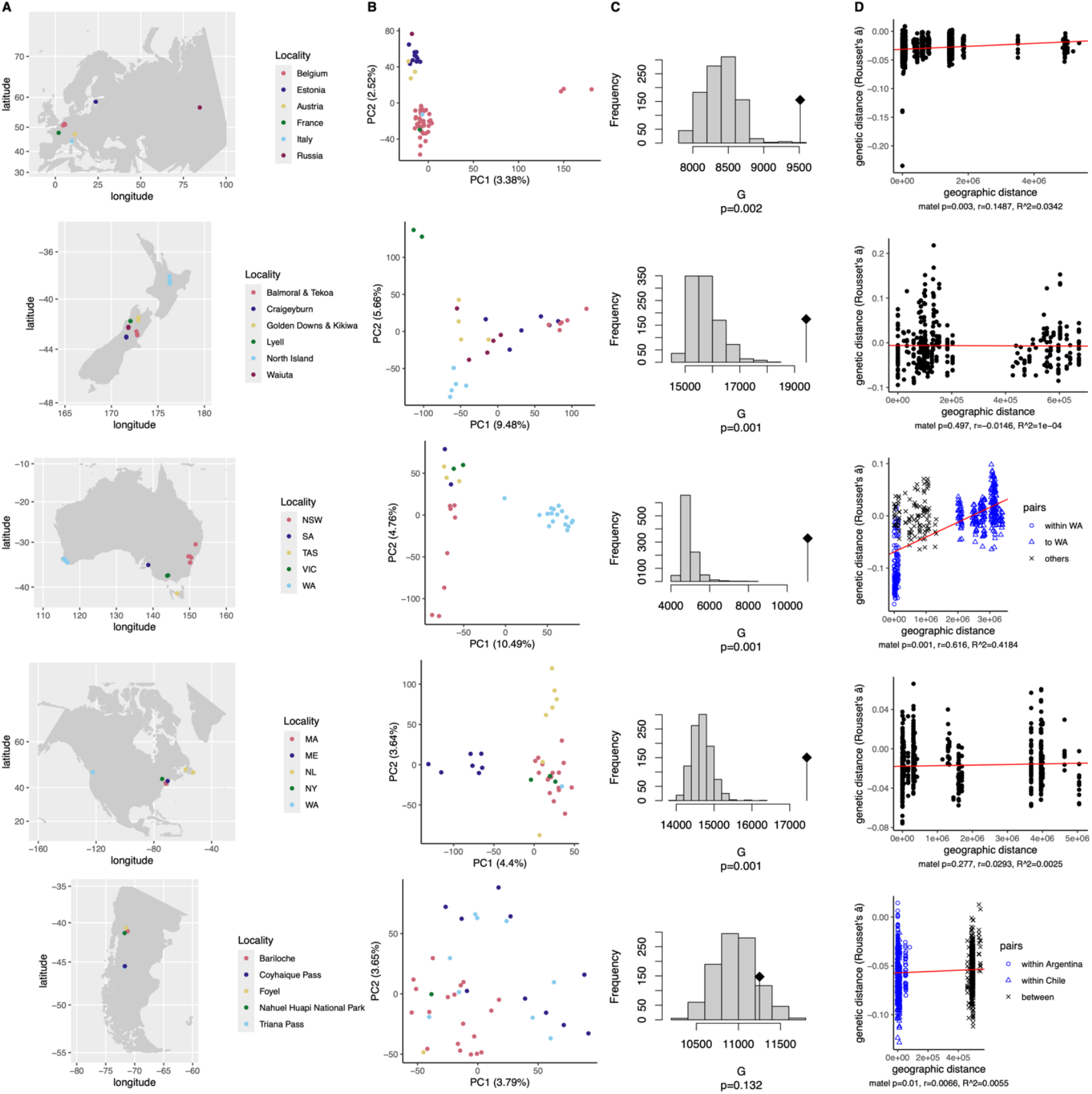
Differentiation of subpopulations within continents in native and introduced *S. luteus* populations. Population structures of individuals collected within each continent were investigated. PCA, G-statistics for the significance of allele frequency difference among localities, and Mantel tests for spatial-genetic relation were applied to reveal subpopulation differentiation within continents. Population structures were detected by G-statistics in all continents except for South America. Among those, the native Europe population had a significant correlation between geographic distance and genetic distance (Mantel test, p<0.05), suggesting a pattern of isolation by distance. For populations in New Zealand and North America, although the subpopulation structures exist (G-test, p=0.001; Fig 6C), the individual-based genetic distance was not correlated to geographic distance (Mantel test p>0.05). In the Australia population, although geographic distance and genetic distance were significantly correlated (Mantel test p=0.001), the pattern was largely driven by the isolated individuals in Western Australia. In South America, since the two populations had no significant differentiation (G-test, p>0.05), the spatial differentiation was in general weak. (A) Maps of sampled subcontinental locations; (B) PCA of subcontinental locations; (C) Distributions of G-statistic by permutation (histograms), observed G-statistic (diamond flag), and p-value of G-statistic; (D) Scatter plot of geographic distance versus genetic distance and p-value of Mantel tests.

No clear isolation by distance pattern was observed within the introduced populations. For populations in New Zealand and North America, although the subpopulation structures exist (G-test, p=0.001; Fig 6C), the individual-based genetic distance was not correlated to geographic distance (Mantel test p>0.05). In the Australia population, although geographic distance and genetic distance were significantly correlated (Mantel test p=0.001), the pattern was largely driven by the isolated individuals in Western Australia. While greater genetic distance was observed between individuals from Western Australia and other parts of Australia (Fig 6 and Fig 3A), the genetic distance of individuals in other parts of Australia did not have a clear correlation to geographic distance (Mantel test p>0.05; Fig S6). Therefore, the pattern was not distinguishable from a single isolated population in Western Australia. In South America, since the two populations had no significant differentiation (G-test, p>0.05), the spatial differentiation was in general weak.

## Discussion

### Human-centered introduction history

Our population genomic analysis showed that all the introduced *S. luteus* populations belong to Clade Central Europe and have little relevance to the populations from Asia or Clade Northern Europe regardless of the introduced geographic localities. This restrictive source of origin, the short time since introduction, plus the significant role several European countries played in the global spread of exotic pine since the 1800s (Vellinga *et al*., 2009; Simberloff *et al*., 2010; Gallien *et al*., 2016) suggests that the introduction of *S. luteus* originated from Europe and was transmitted by human activity instead of natural dispersal. *P. sylvestris*, one of the major native hosts of *S. luteus* and a broadly introduced species across South America, North America, New Zealand, and Australia (Policelli *et al*., 2019), was introduced from Europe. Even though *P. radiata*, the major host pine of *S. luteus* in New Zealand and Australia, is not native to Europe, the pine species was first transplanted to the United Kingdom before being imported to New Zealand and Australia (Shepherd, 1990; Wu *et al*., 2007; Burdon *et al*., 2008). It is plausible that *S. luteus* was introduced from the same route as *P. sylvestris* or *P. radiata* plants as mycorrhizae from Central Europe. Furthermore, given the high genetic diversity and large founder sizes in many introduced populations, the spore bank in the soil carried along with plant seedlings perhaps also played a role in the introductions. Since the inoculum of *S. luteus* was transported in bulk, those transportation routes likely also contain inoculum of other ECM species. Therefore, the same path and process of introduction inferred in *S. luteus* is likely to apply to other species of pine co-introduced ECM fungi as well.

Some of the complex admixture between Australia and New Zealand can also be traced to these countries’ shared history of pine importations and forestry. The establishment of pine plantations in both continents not only involved the introduction of *P. radiata* from the United Kingdom starting from 1857 (Wu *et al*., 2007; Dickie & Smaill, 2019), but also featured exchanges of pine clones between Australia and New Zealand during the breeding process where *P. radiata* was introduced to New Zealand as a seedling transplanted from Australia until 1960s (Shepherd, 1990; Wu *et al*., 2007; Dickie & Smaill, 2019). This historical movement of *P. radiata* aligns with the observed migration from New Zealand to Australia. The sequential introduction of *S. luteus* to Western Australia from other parts of Australia and the strongest population expansion in Western Australia may also be associated with the forestry history. When pine forestry first developed in Western Australia in 1920s, the lack of natural ECM symbionts led to failure in establishing nurseries (Mikola, 1970). The nursery in Hamel, Western Australia ended up introducing soil from other nurseries in Australia as ECM inoculum (Mikola, 1970; Dunstan *et al*., 1998). This nursery later became the major distributor of pine trees in Western Australia. Introducing ECM inoculum artificially from specific sources is likely to result in small numbers of founders, which leads to the smallest initial population size and substantial population expansion in Western Australia. The pattern of sequential introduction suggested by molecular markers agrees with the conjecture that most of pine-symbiotic ECM fungi in Western Australia originated from this single nursery, which acquired ECM inoculum from other parts of Australia. A similar history of artificial introduction of ECM inoculum and a single nursery also explains the severe expansion in Argentina, where the nursery in Isla Victoria was reported as the major nursery from the 1910s to 1960s (Simberloff *et al*., 2010). In addition, ECM inoculum produced by the mycorrhiza laboratory at Casterlar was applied to new nurseries in Argentina (Mikola, 1970). Therefore, as in Western Australia, those two factors of forestry practices are the most plausible causes of the more severe population expansion observed in Argentina.

North American populations underwent little population differentiation and little decrease in genetic diversity. The low levels of evolution and inbreeding since the introduction from Europe suggest that the number of generations, sexual reproduction, and population expansion in North America was limited. Unlike other hosts of *S. luteus* in Australia, New Zealand, and South America, which are typically planted for large-scale forestry, *P. sylvestris*, on which *S. luteus* is dominant in North America, is mainly used for horticultural purposes instead. This relatively limited habitat and host distribution in North America compared to other continents likely limited the expansion and reproduction of introduced *S. luteus*.

### Local genetic variation and structures

The native Central European population exhibits a clear isolation by distance pattern. This suggests the genetic difference is mostly driven by their ability to disperse. This is congruent with the high density and high connectivity of the habitat in Europe. In contrast, in North America, New Zealand, and Australia, the local genetic structures are not driven by physical distance, suggesting migration is not the major factor determining the genetic structure among habitats within those continents. The relatively short time since introductions and fragmented habitats could have prevented the formation of a smooth geographic gradient in their subpopulation structures. Therefore, the structures of the subpopulations are still dominated by the history of human-mediated transport, which may not necessarily correspond to the physical distance between populations. Although *Suillus* spores can be effectively dispersed by air, animals can also contribute to the dispersal of suilloid fungi (Policelli *et al*., 2019). Therefore, the limited interaction with local animal vectors may be another potential reason for impeded dispersal.

Unlike population genetics studies reported in plant pathogens, where distinct clones in introduced populations can be traced back to multiple specific sources (Ali *et al*., 2014; Gladieux *et al*., 2015, 2018; Milgroom *et al*., 2016; Desprez-Loustau *et al*., 2018; Gur *et al*., 2021), the introduced *S. luteus* populations have high genetic diversity and stronger genetic differentiation from the source population and each other. Several factors may have contributed to more signatures of evolution and distinct genetic backgrounds of introduced *S. luteus* populations. First, in comparison to plant pathogens, which experienced strong genetic bottlenecks (Zhou *et al*., 2007; Desprez-Loustau *et al*., 2007; Dutech *et al*., 2012; Gladieux *et al*., 2015, 2018), the larger effective population size of founders observed in *S. luteus* can give rise to more diverse and less clonal genetic backgrounds even within introduced ranges. Second, unlike plant pathogens which frequently reproduce asexually in the introduced range (Goodwin *et al*., 1994; Smart & Fry, 2001; Couch *et al*., 2005; Ali *et al*., 2010; Singh *et al*., 2011; Gladieux *et al*., 2015), the reproduction and dispersal of *S. luteus* and other basidiomycete ECM fungi are predominantly sexual (Coelho *et al*., 2017), which promotes the genetic admixtures in the introduced populations. Third, high levels of outcrossing observed in *Suillus* and other basidiomycete fungi with multi-allelic self-incompatibility systems (Bonello *et al*., 1998; Xu *et al*., 2002; Burchhardt *et al*., 2011; Rivera *et al*., 2014; Kim *et al*., 2015; Branco *et al*., 2017; Zhong *et al*., 2021) are also more likely to preserve genetic variation and repel inbreeding in introduced populations after founder events. Therefore, even though multiple sources of European introductions had occurred, subsequent hybridization over multiple generations may have blurred the boundaries of the sources and created a unique genetic background distinct from the source population.

### *Cryptic species in* Suillus *species*

Phylogenomic analysis revealed previously unknown lineages and expanded the known distribution of *S. luteus* and *S. brunnescens*. Collections of *S. brunnescens* in Japan were observed under native 5-needle pine, *Pinus pumila*, in Hokkaido. This is the first record of *S. brunnescens* outside North America and under *P. pumila*. The deeply divergent clade in combination with the atypical host association and the novel geographic distribution suggests that the Japanese *S. brunnescens* could represent a new species. Notably, its same host association with pines in subgenus *Strobus* as North American *S. brunnescens* (Smith & Thiers, 1964; Nguyen *et al*., 2016) is congruent with the phylogenetic placement, suggesting Japanese *S. brunnescens* has a closer relationship to North American *S. brunnescens* than *S. luteus*, which is instead associated with subgenus *Pinus* (Smith & Thiers, 1964; Nguyen *et al*., 2016). The investigation of morphological differences to elevate Japanese *S. brunnescens* to a new species requires additional study.

Two newly discovered divergent clades of *S. luteus* were detected in this study, from Asia and Northern Europe. Their phylogeny, PCA placement, admixture analysis, and genetic distances all suggest more distant relationships and limited gene flow between those two clades and Clade Central Europe. Some collections of Clade Northern Europe and Clade Central Europe were found in geographically proximate locations, including collections in two adjacent provinces in Russia and collections within Austria, both under the same species of pine host. In these cases, the differentiation strongly implies sympatric reproductive isolation as cryptic species instead of population differentiation due to geographic or host-specificity barriers. This further suggests that Clade Asia, which is sister to Clade Northern Europe, also represents a distinct species from Clade Central Europe. Whether morphological differences exist among those clades still requires further observations from additional collections.

### Conclusion

This study provides the first detailed global population analysis of *S. luteus* and the inference of the introduction process of pine-co-introduced ECM fungi at the population level. It highlights that while the European origin is consistent for all introduced populations, the patterns of subsequent admixtures, genetic differentiation, and population expansion vary among different introduced populations. With these results, we are able to link the history of human-contributed movement of the fungal species to forestry practices. The absence of isolation by distance in the introduced populations highlights the disparity in the dispersal ecology and history of *S. luteus* populations between its native and exotic ranges. Lastly, potential lineages of cryptic species indicate the unexplored biodiversity within *S. luteus* and other *Suillus* species whose collections have not been studied across wide geographic ranges. Although our focus lies on *S. luteus*, the observed population genetic patterns are likely also applicable to other pine-co-introduced ECM fungi that were transported via the same route as *S. luteus*.

## Acknowledgments

This study would not have been possible without the contributions of collaborators who provided specimens, cultures, and who facilitated fieldwork for this study. We are grateful to all of them for their assistance, especially Jeff Powell, Susan Hopkins, Amanda Certano, Nahuel Policelli, Martin Nunez, Jonathan Frank, Jessie Uehling, Audrius Menkis, Tatiana Buyonkova, Nina Filippova, Nadezhda Agaphonova, Otto Miettinen, Fred Asiegbu, Jan Colpert, Tartarotti Siegfried, Andrus Voitk, Yolanda Wiersma, Linas Kudzma, Teresa Lebel, Nimal Karunajeewa, Regina Kuhnert, Tom May, Pamela and David Catcheside, Neale Bougher, Mark Brundrett, Bill Dunstan, Andrew Bissett, Tim Wardlaw, Peter Thrall, Pat Leonard, Maj Padamsee, Peter Buchanan, Peter Johnston, Sarah Addison, Ian Hood, and Yiyin Chang.

We thank the staff and the researchers at DOE Joint Genome Institute who handled and managed DNA samples and sequencing data. We thank the Office of Information Technology and Research Computing team at Duke University for maintaining and providing computational resources.

The work (proposal: doi.org/10.46936/10.25585/60007225) conducted by the U.S. Department of Energy Joint Genome Institute (https://ror.org/04xm1d337), a DOE Office of Science User Facility, is supported by the Office of Science of the U.S. Department of Energy operated under Contract No. DE-AC02-05CH11231. Financial support for this work was provided through National Science Foundation Grants DEB 1554181 to R. Vilgalys and P. Kennedy, IOS-PBI 2029168 to S. Branco and H.-L. Liao, OISE 1953299 to J. Hoeksema, and Research Foundation Flanders grant FWO-G082612N to J. Ruytinx. L. Lofgren is supported by funding from the National Institutes of Health grant no. T32-AI052080 via the Tri-I MMPTP Fellowship.

## Author contributions

YHK, RV, JH, NHN, JR, TB, SB, PK, IG, JAR, and HLL developed the scope and concept of the work. YHK, RV, JR, TB, UP, JP, LT, PK, LB, and AR identified the sampling sites and collected the samples. YHK processed the samples. YHK directed and conducted the analyses. AB, SB, LL and AR assisted in formulating the analyses. AK, AL, and KB managed and processed the sequencing data. YHK and RV prepared the manuscript. All authors contributed with edits and suggestions for the analyses and manuscript.

## Data Availability

The genome sequencing data was deposited to NCBI SRA available under BioProject PRJNA1029143. The accession numbers of each sample are listed in Supplementary Table S2. The scripts used for analysis are available on GitHub (https://github.com/KeFungi/Suilu_PopStructure).

## References

Alexander DH, Novembre J, Lange K. 2009. Fast model-based estimation of ancestry in unrelated individuals. Genome Research 19: 1655–64.

Ali S, Gladieux P, Leconte M, Gautier A, Justesen AF, Hovmøller MS, Enjalbert J, Vallavieille-Pope C de. 2014. Origin, Migration Routes and Worldwide Population Genetic Structure of the Wheat Yellow Rust Pathogen Puccinia striiformis f.sp. tritici. PLOS Pathogens 10: e1003903.

Ali S, Leconte M, Walker A-S, Enjalbert J, de Vallavieille-Pope C. 2010. Reduction in the sex ability of worldwide clonal populations of Puccinia striiformis f.sp. tritici. Fungal Genetics and Biology 47: 828–838.

van der Auwera GA, O’Connor BD. 2020. Genomics in the Cloud: using Docker, GATK, and WDL in Terra. O’Reilly Media.

Bazzicalupo AL, Ruytinx J, Ke YH, Coninx L, Colpaert JV, Nguyen NH, Vilgalys R, Branco S. 2020. Fungal heavy metal adaptation through single nucleotide polymorphisms and copy-number variation. Molecular Ecology 29: 4157–4169.

Bonello P, Bruns TD, Gardes M. 1998. Genetic structure of a natural population of the ectomycorrhizal fungus Suillus pungens. New Phytologist 138: 533–542.

Branco S, Bi K, Liao H-L, Gladieux P, Badouin H, Ellison CE, Nguyen NH, Vilgalys R, Peay KG, Taylor JW, et al. 2017. Continental-level population differentiation and environmental adaptation in the mushroom Suillus brevipes. Molecular Ecology 26: 2063–2076.

Branco S, Gladieux P, Ellison CE, Kuo A, LaButti K, Lipzen A, Grigoriev IV, Liao HL, Vilgalys R, Peay KG, et al. 2015. Genetic isolation between two recently diverged populations of a symbiotic fungus. Molecular Ecology 24: 2747–58.

Burchhardt KM, Rivera Y, Baldwin T, VanEarden D, Kretzer AM. 2011. Analysis of genet size and local gene flow in the ectomycorrhizal basidiomycete Suillus spraguei (synonym S. pictus). Mycologia 103: 722–730.

Burdon RD, Carson MJ, Shelbourne CJA. 2008. Achievements in forest tree genetic improvement in Australia and New Zealand 10: Pinus radiata in New Zealand. Australian Forestry 71: 263–279.

Chang CC, Chow CC, Tellier LC, Vattikuti S, Purcell SM, Lee JJ. 2015. Second-generation PLINK: rising to the challenge of larger and richer datasets. Gigascience 4: 7.

Chapela IH, Osher LJ, Horton TR, Henn MR. 2001. Ectomycorrhizal fungi introduced with exotic pine plantations induce soil carbon depletion. Soil Biology and Biochemistry 33: 1733–1740.

Coelho MA, Bakkeren G, Sun S, Hood ME, Giraud T. 2017. Fungal sex: the Basidiomycota. Microbiology Spectrum 5.

Couch BC, Fudal I, Lebrun MH, Tharreau D, Valent B, van Kim P, Notteghem JL, Kohn LM. 2005. Origins of host-specific populations of the blast pathogen Magnaporthe oryzae in crop domestication with subsequent expansion of pandemic clones on rice and weeds of rice. Genetics 170: 613–30.

Desprez-Loustau M-L, Massot M, Toïgo M, Fort T, Aday Kaya AG, Boberg J, Braun U, Capdevielle X, Cech T, Chandelier A, et al. 2018. From leaf to continent: The multi-scale distribution of an invasive cryptic pathogen complex on oak. Fungal Ecology 36: 39–50.

Desprez-Loustau M-L, Robin C, Buée M, Courtecuisse R, Garbaye J, Suffert F, Sache I, Rizzo DM. 2007. The fungal dimension of biological invasions. Trends in Ecology & Evolution 22: 472–480.

Dickie IA, Bolstridge N, Cooper JA, Peltzer DA. 2010. Co-invasion by Pinus and its mycorrhizal fungi. New Phytologist 187: 475–484.

Dickie IA, Bufford JL, Cobb RC, Desprez-Loustau M-L, Grelet G, Hulme PE, Klironomos J, Makiola A, Nuñez MA, Pringle A, et al. 2017. The emerging science of linked plant–fungal invasions. New Phytologist 215: 1314–1332.

Dickie IA, Smaill SJ. 2019. The role of mycorrhizal fungi in plant invasions. Canterbury Botanical Society Journal 50.

Douhan GW, Vincenot L, Gryta H, Selosse M-A. 2011. Population genetics of ectomycorrhizal fungi: from current knowledge to emerging directions. Fungal Biology 115: 569–597.

Dray S, Dufour A-B. 2007. The ade4 package: implementing the duality diagram for ecologists. Journal of Statistical Software 22: 1–20.

Dunstan WA, Dell B, Malajczuk N. 1998. The diversity of ectomycorrhizal fungi associated with introduced Pinus spp. in the Southern Hemisphere, with particular reference to Western Australia. Mycorrhiza 8: 71–79.

Dutech C, Barrès B, Bridier J, Robin C, Milgroom MG, Ravigné V. 2012. The chestnut blight fungus world tour: successive introduction events from diverse origins in an invasive plant fungal pathogen. Molecular Ecology 21: 3931–3946.

Gallien L, Saladin B, Boucher FC, Richardson DM, Zimmermann NE. 2016. Does the legacy of historical biogeography shape current invasiveness in pines? New Phytologist 209: 1096–1105.

Gladieux P, Feurtey A, Hood ME, Snirc A, Clavel J, Dutech C, Roy M, Giraud T. 2015. The population biology of fungal invasions. Molecular Ecology 24: 1969–1986.

Gladieux P, Ravel S, Rieux A, Cros-Arteil S, Adreit H, Milazzo J, Thierry M, Fournier E, Terauchi R, Tharreau D. 2018. Coexistence of multiple endemic and pandemic lineages of the rice blast pathogen. mBio 9: e01806–17.

Goodwin SB, Cohen BA, Fry WE. 1994. Panglobal distribution of a single clonal lineage of the Irish potato famine fungus. Proceedings of the National Academy of Sciences 91: 11591–11595.

Goudet J. 2005. hierfstat, a package for R to compute and test hierarchical F-statistics. Molecular Ecology Notes 5: 184–186.

Goudet J, Raymond M, de Meeus T, Rousset F. 1996. Testing differentiation in diploid populations. Genetics 144: 1933–40.

Gur L, Reuveni M, Cohen Y, Cadle-Davidson L, Kisselstein B, Ovadia S, Frenkel O. 2021. Population structure of Erysiphe necator on domesticated and wild vines in the Middle East raises questions on the origin of the grapevine powdery mildew pathogen. Environmental Microbiology 23.

van der Heijden MGA, Martin FM, Selosse M-A, Sanders IR. 2015. Mycorrhizal ecology and evolution: the past, the present, and the future. New Phytologist 205: 1406–1423.

Hoeksema JD, Averill C, Bhatnagar JM, Brzostek E, Buscardo E, Chen K-H, Liao H-L, Nagy L, Policelli N, Ridgeway J, et al. 2020. Ectomycorrhizal plant-fungal co-invasions as natural experiments for connecting plant and fungal traits to their ecosystem consequences. Frontiers in Forests and Global Change 3.

Houtgast EJ, Sima VM, Bertels K, Al-Ars Z. 2018. Hardware acceleration of BWA-MEM genomic short read mapping for longer read lengths. Computational Biology and Chemistry 75: 54–64.

Huson DH, Bryant D. 2006. Application of phylogenetic networks in evolutionary studies. Molecular Biology and Evolution 23: 254–267.

Jombart T, Ahmed I. 2011. adegenet 1. 3-1: new tools for the analysis of genome-wide SNP data. Bioinformatics 27: 3070–1.

Kamm J, Terhorst J, Durbin R, Song YS. 2020. Efficiently inferring the demographic history of many populations with allele count data. Journal of the American Statistical Association 115: 1472–1487.

Kim KH, Ka KH, Kang JH, Kim S, Lee JW, Jeon BK, Yun JK, Park SR, Lee HJ. 2015. Identification of single nucleotide polymorphism markers in the laccase gene of shiitake mushrooms (Lentinula edodes). Mycobiology 43: 75–80.

Kohler A, Kuo A, Nagy LG, Morin E, Barry KW, Buscot F, Canback B, Choi C, Cichocki N, Clum A, et al. 2015. Convergent losses of decay mechanisms and rapid turnover of symbiosis genes in mycorrhizal mutualists. Nature Genetics 47: 410–5.

Manichaikul A, Mychaleckyj JC, Rich SS, Daly K, Sale M, Chen WM. 2010. Robust relationship inference in genome-wide association studies. Bioinformatics 26: 2867–73.

Martin M. 2011. Cutadapt removes adapter sequences from high-throughput sequencing reads. EMBnet.journal 17: 10.

Marx DH. 1969. The influence of ectotrophic mycorrhizal fungi on the resistance of pine roots to pathogenic infections. II. Production, identification, and biological activity of antibiotics produced by Leucopaxillus cerealis var. piceina. Phytopathology 59: 411–7.

Meyerson LA, Mooney HA. 2007. Invasive alien species in an era of globalization. Frontiers in Ecology and the Environment 5: 199–208.

Mikola P. 1970. Mycorrhizal inoculation in afforestation. In: Romberger JA, Mikola P, eds. International Review of Forestry Research. International Review of Forestry Research. Elsevier, 123–196.

Milgroom MG, del Mar Jiménez-Gasco M, Olivares-García C, Jiménez-Díaz RM. 2016. Clonal Expansion and Migration of a Highly Virulent, Defoliating Lineage of Verticillium dahliae. Phytopathology 106: 1038–1046.

Mirov NT. 1967. The genus Pinus. New York: Ronald Press Company.

Nguyen NH, Vellinga EC, Bruns TD, Kennedy PG. 2016. Phylogenetic assessment of global Suillus ITS sequences supports morphologically defined species and reveals synonymous and undescribed taxa. Mycologia 108: 1216–1228.

Nuñez MA, Dickie IA. 2014. Invasive belowground mutualists of woody plants. Biological Invasions 16: 645–661.

Peck CH. 1887. New York species of viscid boleti. Bulletin of the New York State Museum 1: 57–66.

Policelli N, Bruns TD, Vilgalys R, Nuñez MA. 2019. Suilloid fungi as global drivers of pine invasions. New Phytologist 222: 714–725.

Policelli N, Hoeksema JD, Moyano J, Vilgalys R, Vivelo S, Bhatnagar JM. 2023. Global pine tree invasions are linked to invasive root symbionts. New Phytologist 237: 16–21.

Policelli N, Nuñez MA. 2025. Invasive ectomycorrhizal fungi: belowground insights from South America. New Phytologist 248: 2714–2721.

Pringle A, Bever JD, Gardes M, Parrent JL, Rillig MC, Klironomos JN. 2009. Mycorrhizal symbioses and plant invasions. Annual Review of Ecology, Evolution, and Systematics 40: 699–715.

Rejmánek M, Richardson DM. 2013. Trees and shrubs as invasive alien species – 2013 update of the global database. Diversity and Distributions 19: 1093–1094.

Reynolds J, Weir BS, Cockerham CC. 1983. Estimation of the coancestry coefficient: basis for a short-term genetic distance. Genetics 105: 767–79.

Rivera Y, Burchhardt KM, Kretzer AM. 2014. Little to no genetic structure in the ectomycorrhizal basidiomycete Suillus spraguei (Syn. S. pictus) across parts of the Northeastern USA. Mycorrhiza 24: 227–32.

Rousset F. 2008. genepop’007: a complete re-implementation of the genepop software for Windows and Linux. Molecular Ecology Resources 8: 103–106.

Rousset, F. 2000. Genetic differentiation between individuals. Journal of Evolutionary Biology 13: 58–62.

Sakai AK, Allendorf FW, Holt JS, Lodge DM, Molofsky J, With KA, Baughman S, Cabin RJ, Cohen JE, Ellstrand NC, et al. 2001. The population biology of invasive species. Annual Review of Ecology and Systematics 32: 305–332.

Selosse M-A, Baudoin E, Vandenkoornhuyse P. 2004. Symbiotic microorganisms, a key for ecological success and protection of plants. Comptes Rendus Biologies 327: 639–648.

Shaw PJA, Kibby C, Mayes J. 2003. Effects of thinning treatment on an ectomycorrhizal succession under Scots pine. Mycological Research 107: 317–328.

Shepherd RW. 1990. Early importations of Pinus radiata to New Zealand and distribution in Canterbury to 1885: implications for the genetic makeup of Pinus radiata stocks. Part I. Horticulture in New Zealand 1: 33–38.

Simberloff D, Nuñez MA, Ledgard NJ, Pauchard A, Richardson DM, Sarasola M, Van Wilgen BW, Zalba SM, Zenni RD, Bustamante R, et al. 2010. Spread and impact of introduced conifers in South America: Lessons from other southern hemisphere regions. Austral Ecology 35: 489–504.

Singh RP, Hodson DP, Huerta-Espino J, Jin Y, Bhavani S, Njau P, Herrera-Foessel S, Singh PK, Singh S, Govindan V. 2011. The Emergence of Ug99 Races of the Stem Rust Fungus is a Threat to World Wheat Production. Annual Review of Phytopathology 49: 465–481.

Skilling DD. 1990. Pinus sylvestris L. Scotch pine. Silvics of North America 11: 489–496.

Skrede I, Murat C, Hess J, Maurice S, Sonstebo JH, Kohler A, Barry-Etienne D, Eastwood D, Hogberg N, Martin F, et al. 2021. Contrasting demographic histories revealed in two invasive populations of the dry rot fungus Serpula lacrymans. Molecular Ecology 30: 2772–2789.

Smart CD, Fry WE. 2001. Invasions by the Late Blight Pathogen: Renewed Sex and Enhanced Fitness. Biological Invasions 3: 235–243.

Smith SE, Read DJ. 2010. Mycorrhizal symbiosis. Academic press.

Smith AH, Thiers HD. 1964. A contribution toward a monograph of North American species of Suillus. Ann Arbor: Privately published.

Stamatakis A. 2014. RAxML version 8: a tool for phylogenetic analysis and post-analysis of large phylogenies. Bioinformatics 30: 1312–3.

Strullu-Derrien C, Selosse M-A, Kenrick P, Martin FM. 2018. The origin and evolution of mycorrhizal symbioses: from palaeomycology to phylogenomics. New Phytologist 220: 1012–1030.

Tajima F. 1989. Statistical method for testing the neutral mutation hypothesis by DNA polymorphism. Genetics 123: 585–95.

Tedersoo L, Brundrett MC. 2017. Evolution of Ectomycorrhizal Symbiosis in Plants. In: Tedersoo L, ed. Biogeography of Mycorrhizal Symbiosis. Switzerland: Springer International Publishing, 407–467.

Tedersoo L, Suvi T, Beaver K, Kõljalg U. 2007. Ectomycorrhizal fungi of the Seychelles: diversity patterns and host shifts from the native Vateriopsis seychellarum (Dipterocarpaceae) and Intsia bijuga (Caesalpiniaceae) to the introduced Eucalyptus robusta (Myrtaceae), but not Pinus caribea (Pinaceae). New Phytologist 175: 321–333.

Vellinga EC, Wolfe BE, Pringle A. 2009. Global patterns of ectomycorrhizal introductions. New Phytologist 181: 960–973.

Vietorisz CR, Nash JA, Siggers JA, Leander EJ, Bock BM, Camuy-Vélez LA, Hall AJ, Jaros JE, Kuehn KA, Lai EY, et al. 2025. Pine-fungal co-invasion alters whole-ecosystem properties of a native eucalypt forest. New Phytologist 247: 2342–2356.

Westphal MI, Browne M, MacKinnon K, Noble I. 2008. The link between international trade and the global distribution of invasive alien species. Biological Invasions 10: 391–398.

Wu HX, Eldridge KG, Matheson AColin, Powell MB, McRae TA, Butcher TB, Johnson IG. 2007. Achievements in forest tree improvement in Australia and New Zealand 8. Successful introduction and breeding of radiata pine in Australia. Australian Forestry 70: 215–225.

Xu J, Desmerger C, Callac P. 2002. Fine-scale genetic analyses reveal unexpected spatial-temporal heterogeneity in two natural populations of the commercial mushroom Agaricus bisporus. Microbiology 148: 1253–1262.

Zhong J, Xu J, Zhang P. 2021. Diversity, dispersal and mode of reproduction of Amanita exitialis in southern china. Genes 12: 1907.

Zhou X, Burgess TI, De Beer ZW, Lieutier F, Yart A, Klepzig K, Carnegie A, Portales JM, Wingfield BD, Wingfield MJ. 2007. High intercontinental migration rates and population admixture in the sapstain fungus Ophiostoma ips. Molecular Ecology 16: 89–99.

