## supplemental figures for "Global population structures and demographic history of *Suillus luteus*, a pine co-introduced ectomycorrhizal fungus associated with exotic forestry and invasion"

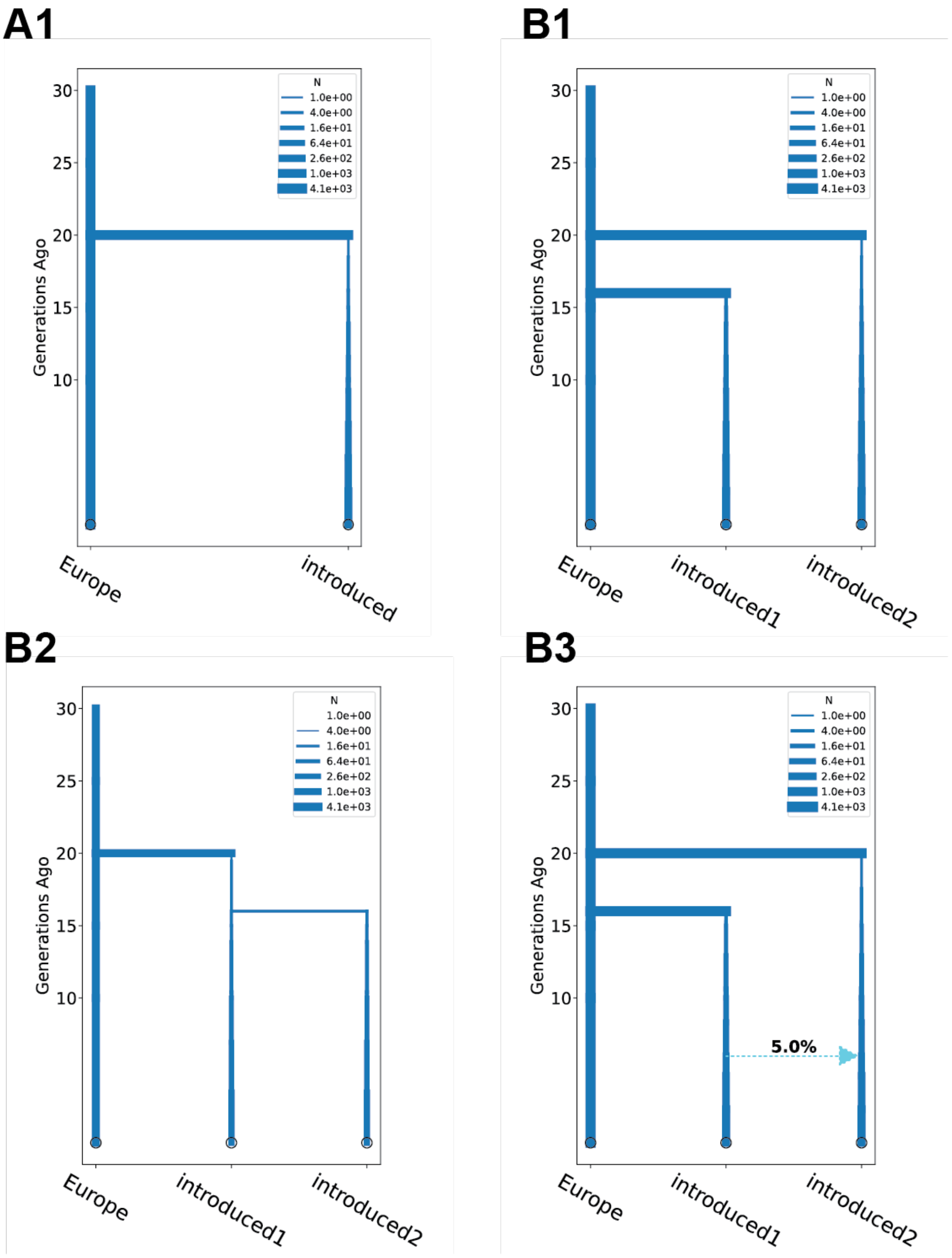
A1
B1
B2
B3
N
1.0e+00
4.0e+00
1.6e+01
6.4e+01
2.6e+02
1.0e+03
4.1e+03
Generations Ago
30
25
20
15
10
Europe
introduced
introduced1
introduced2
5.0%

**Figure S1 Diagrams of divergence histories considered in demographic modeling.**

Demographic models used in SFS-based inference in momi2. **A1** contains divergence with a constant growth rate. **B1** is designed to fit the scenario of independent introductions without subsequent admixture. Two introduced populations independently diverged from the Europe population some time ago. **B2** is designed to fit the scenario of a sequential introduction. One introduced population diverges from the Europe population followed by the other population diverging from the first introduced population. **B3** is designed to fit the scenario of independent introductions with admixture. Two introduced populations independently diverge from the Europe population, followed by a single pulse of unidirectional migration. Constant growth rates are considered in all B1-B3. Divergence time, population size, and growth rate are set as independent parameters.

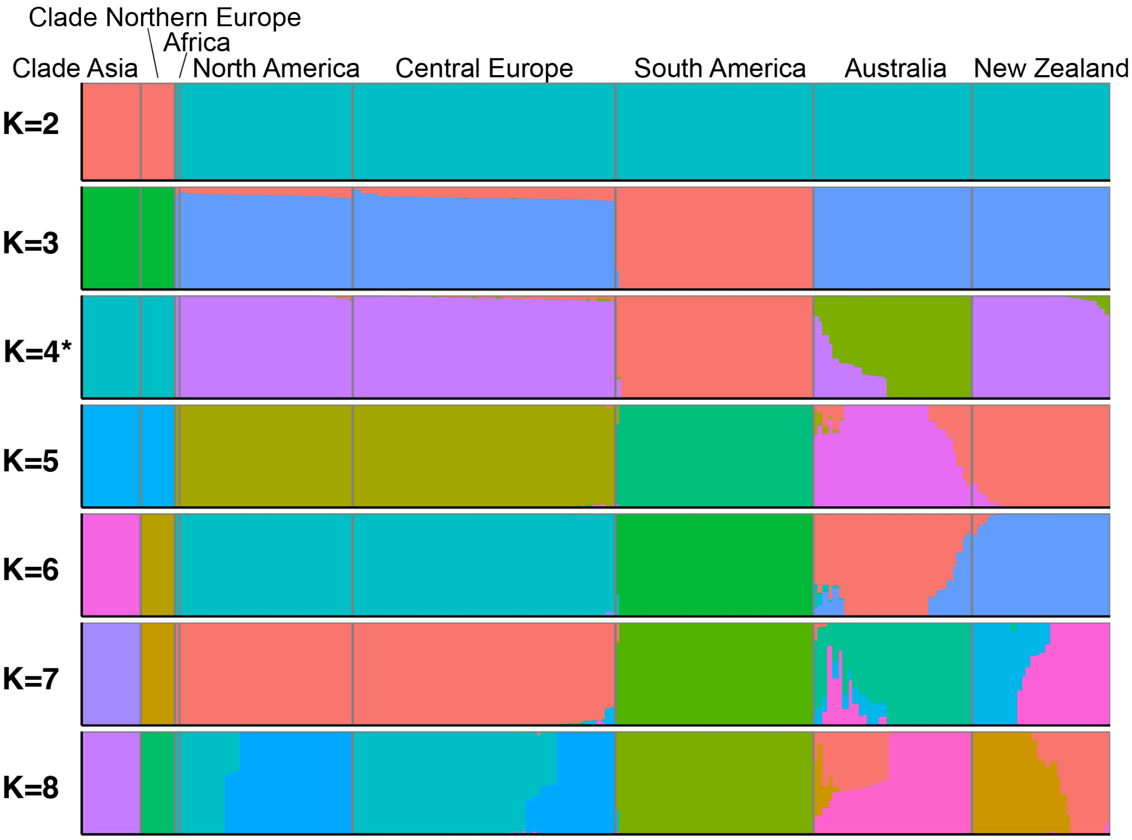


**Figure S2 Ancestry of individuals among global populations of *Suillus luteus*.**

Admixture analysis of *S. luteus* individuals from all clades inferred from 453,945 LD-pruned SNPs without prior population assignment. K=4 was determined to be the optimal number of population clusters based on the criteria of cross-validation. The optimal clusters suggest the individuals in Clade Asia and Clade Northern Europe had distinct ancestry from populations in Clade Central Europe without gene flow. Among populations in Clade Central Europe, South America had the highest level of differentiation from the Central Europe population and little evidence of admixture. Differences between North America and Europe were not detectable. Evidence of admixture was found between populations in Australia and New Zealand.

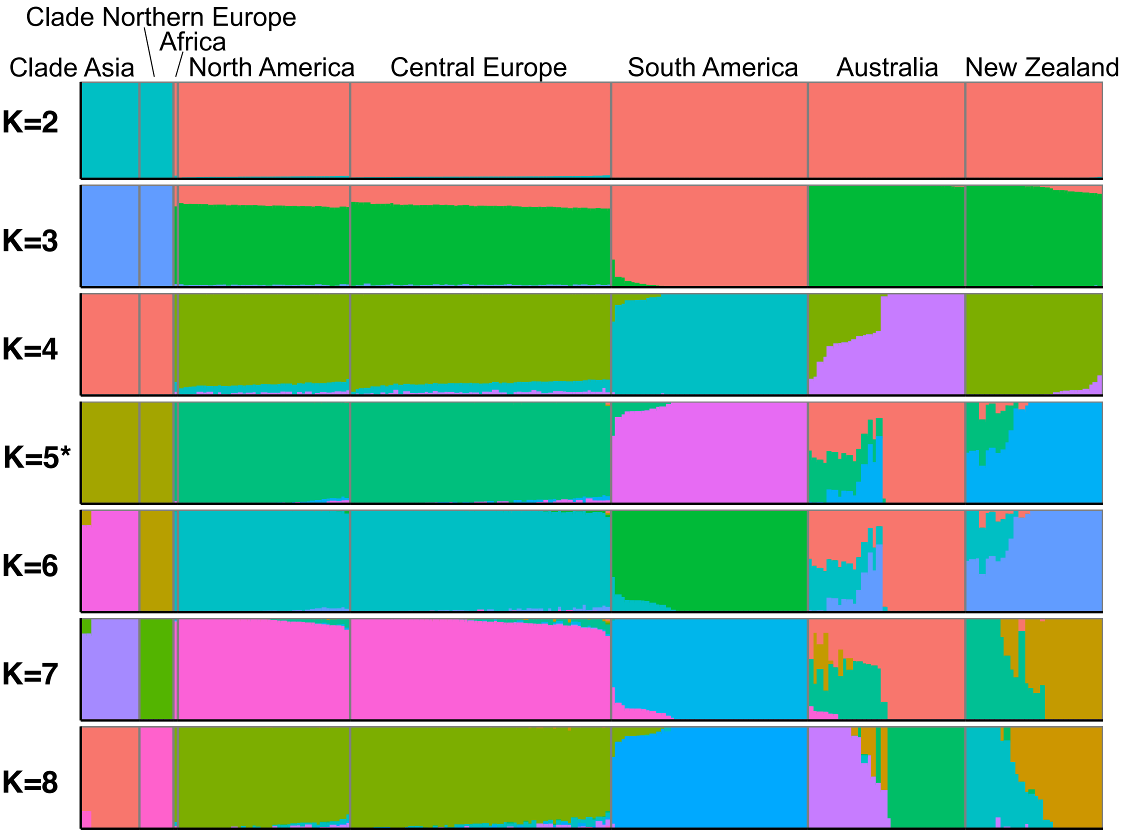


**Figure S3 Ancestry of individuals among global populations of *Suillus luteus* with SNPs having minor allele frequency greater than 0.025.**
The same admixture analysis *of S. luteus* individuals from all clades as Figure S2, except inferred from 138,697 SNPs whose minor allele frequencies across all populations were greater than 0.025.

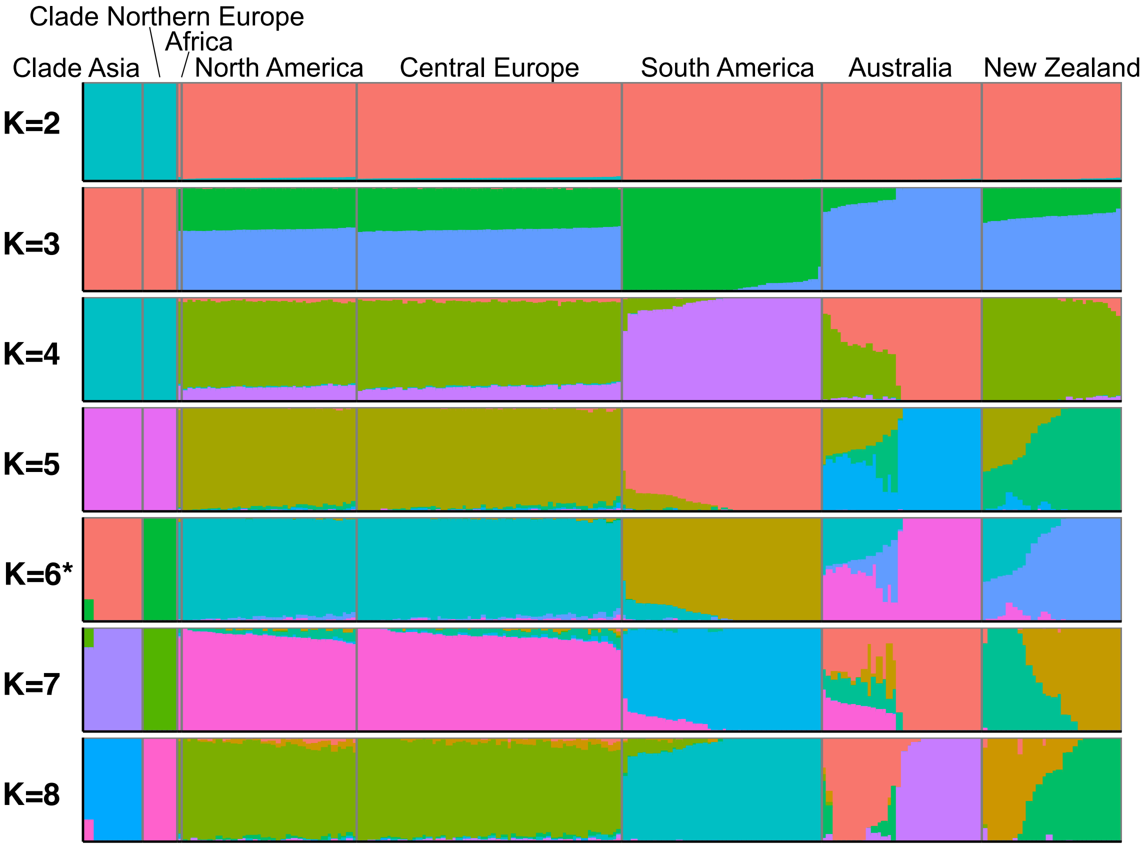


**Figure S4 Ancestry of individuals among global populations of *Suillus luteus* with SNPs having minor allele frequency greater than 0.05.**

The same admixture analysis *of S. luteus* individuals from all clades as Figure S2, except inferred from 80,032 SNPs whose minor allele frequencies across all populations were greater than 0.05.

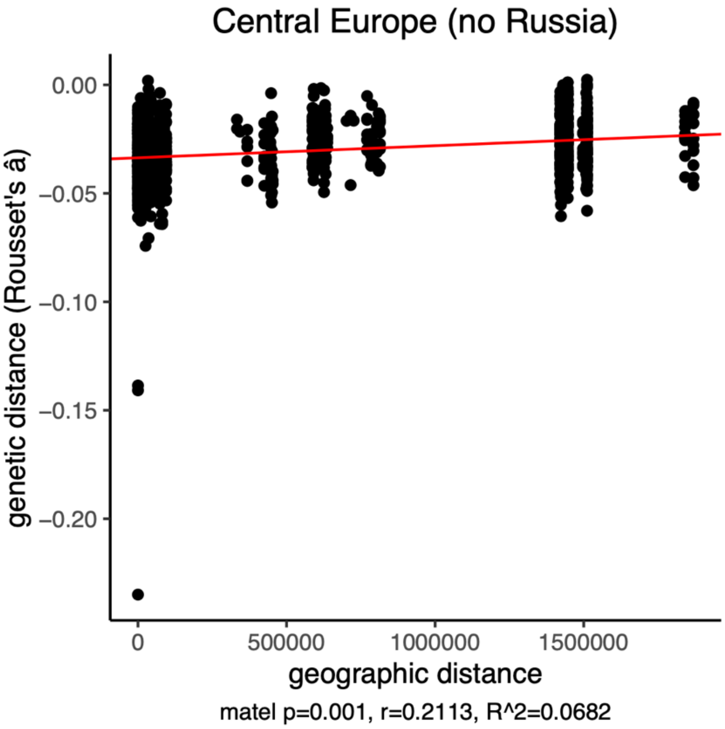


**Figure S5. Scatter plot of geographic distance versus genetic distance, and Mantel tests p-value for Central Europe excluding individuals from Russia.**

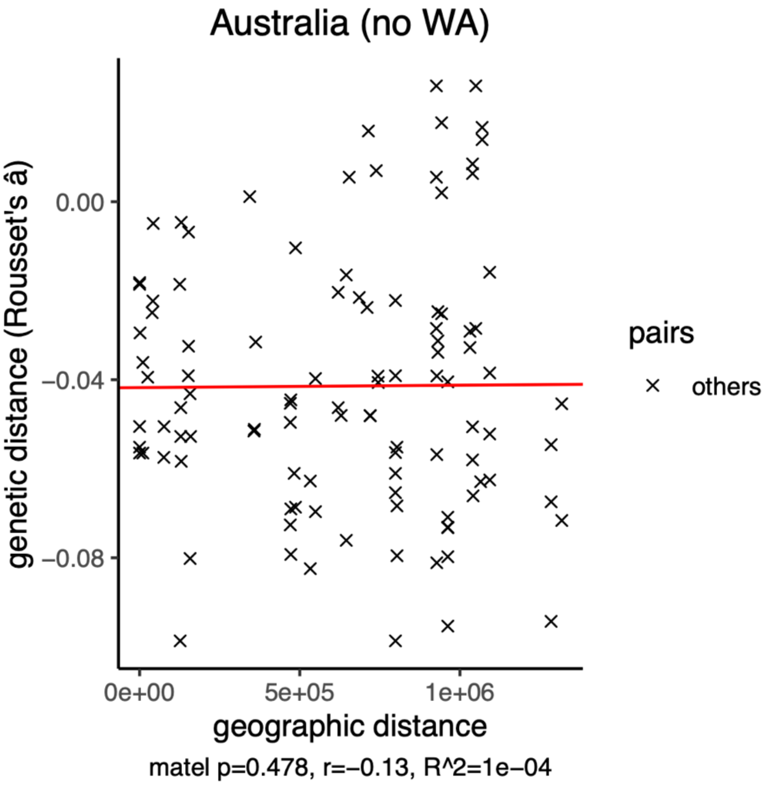


**Figure S6. Scatter plot of geographic distance versus genetic distance, and Mantel tests p-value for Australia excluding individuals from Western Australia.**
