## supplemental methods for "Global population structures and demographic history of *Suillus luteus*, a pine co-introduced ectomycorrhizal fungus associated with exotic forestry and invasion"

### Supplementary Methods
#### Individuals excluded from analysis
Some genomes sequenced and available for this project were not included in the final analysis. The reasons and criteria are described below.

*Haploid genome*

We observed that some genomes have practically zero heterozygous sites. Their percent of heterozygous sites is disjunct from other individuals in the same population (Figure M1). The calculation of heterozygous sites was not attributed to bias due to their dissimilarity to the reference genome, since genomes of other species (*S. brevipes* and *S. brunnescens*) did not fail this criterion. Notably, all genomes belonging to this category were sequenced from cultures, suggesting dedikaryotization may have occurred during serial transfer of cultures and/or selection by culture media (Ginterova, 1973; Arita, 1979; Fukumasa-Nakai *et al.*, 1994). Among those few sites detected as heterozygous, the alternative allele frequency (reference/alternative assigned randomly) deviated from 0.5 and is polymodal (black plots in Figure M2), implying the lack of genuine heterozygous sites except noise signals. Last, those genomes all missed any heterozygous sites around HD MAT mating type locus (Figure M3), indicating those genomes only possessed a single mating type. Therefore, we excluded those *S. luteus* genomes of the number of heterozygous sites below a hard cutoff 2% as shown in Figure M1. Culture aurim734, a long-term culture isolated from wild pine root with only one mating type and lower heterozygous sites, was also excluded.

*Mixed collection/contamination*

Some genomes have visible bimodal distribution of alternative allele frequency (red plots in Figure M2), which indicates the heterozygous sites are mostly from a mixture of individuals instead of from a single diploid genome. This is likely the result of mixed collection or cross-contaminating genomic DNA. Therefore, these individuals were excluded from analyses.

*Clonal and closely related individuals*

We endeavored to collect samples over 10 meters apart to avoid duplicated sampling of the same genet. However, some genets can colonize as far as 16-30 meters in diameter in *Suillus* spp. (Douhan *et al.*, 2011). The Hamming distance between each pair of individuals from the same population was calculated by command *--distance square 1-ibs* in *plink 1.9* (Chang *et al.*, 2015). Near zero genetic distance were observed between several pair of genomes in the same location (NZ006:NZ008, NZ169:NZ75, UH-Slu-LM12:UH-Slu-LM5, DOB636:DOB637, AU463:AU464, UH-Slu-Sl22:UH-Slu-Sl24, JS063:JS064, SUI20:SUI22:SUI23, and TW03:TW12) (Figure M4). Therefore, given the high likelihoods that they belong to the same genet, cutoff value 0.01 was applied to exclude collections from the same individual. Only the first collection (lower collection number) was included in the analyses. Three individuals collected in Europe from the same location, UH-Slu-Ew1, UH-Slu-Ew2, and UH-Slu-Ew7, also had a high kinship coefficient (> 0.2). Those samples were excluded from analyses that are sensitive to inbreeding and the increase of rare alleles, namely Tajima's D and SFS-based modeling.

*Cultures from spore prints*

Cultures MS-1-16, MS-1-6, MS-3-17, MS-5-18, MS-5-39, MS-5-42, and MS-5-44 were isolated from pine root tips inoculated by single spore print. Those dikaryotic cultures became offspring of selfing by their nature. This would bias the parameter estimations assuming random mating and affect the estimation of selfing rate. Moreover, there is possibility of crossing spores from individuals in different populations due to unintentional cross-contamination. Taking those concerns together, those samples were also excluded from downstream analyses.

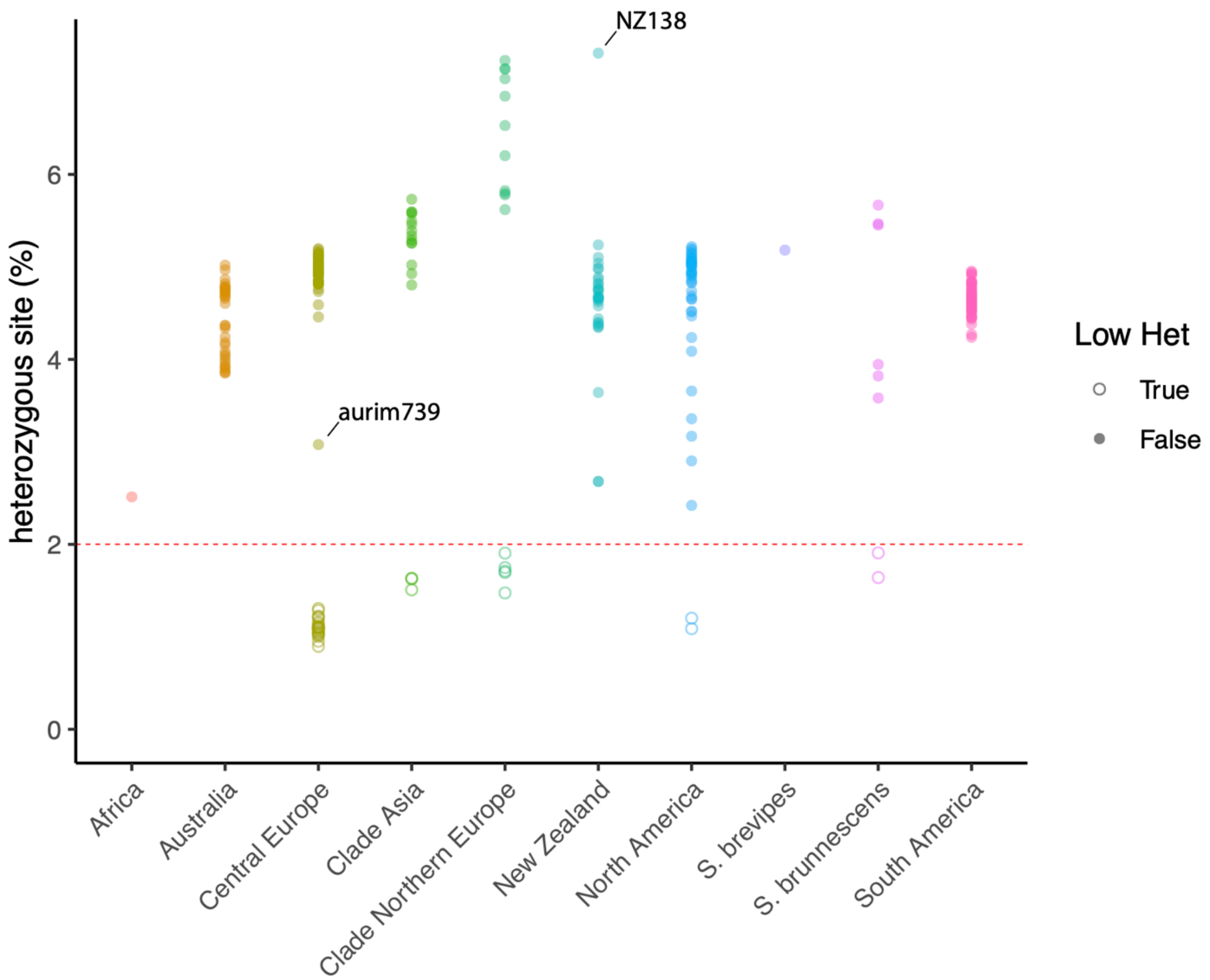


**Figure M1. Percentage of heterozygous sites of each genome.**
Genomes with less than 2% heterozygous sites (hollow circles) were excluded from analyses. Culture aurim734, a long-term culture isolated from wild pine root with only one mating type and lower heterozygous site, was also excluded.

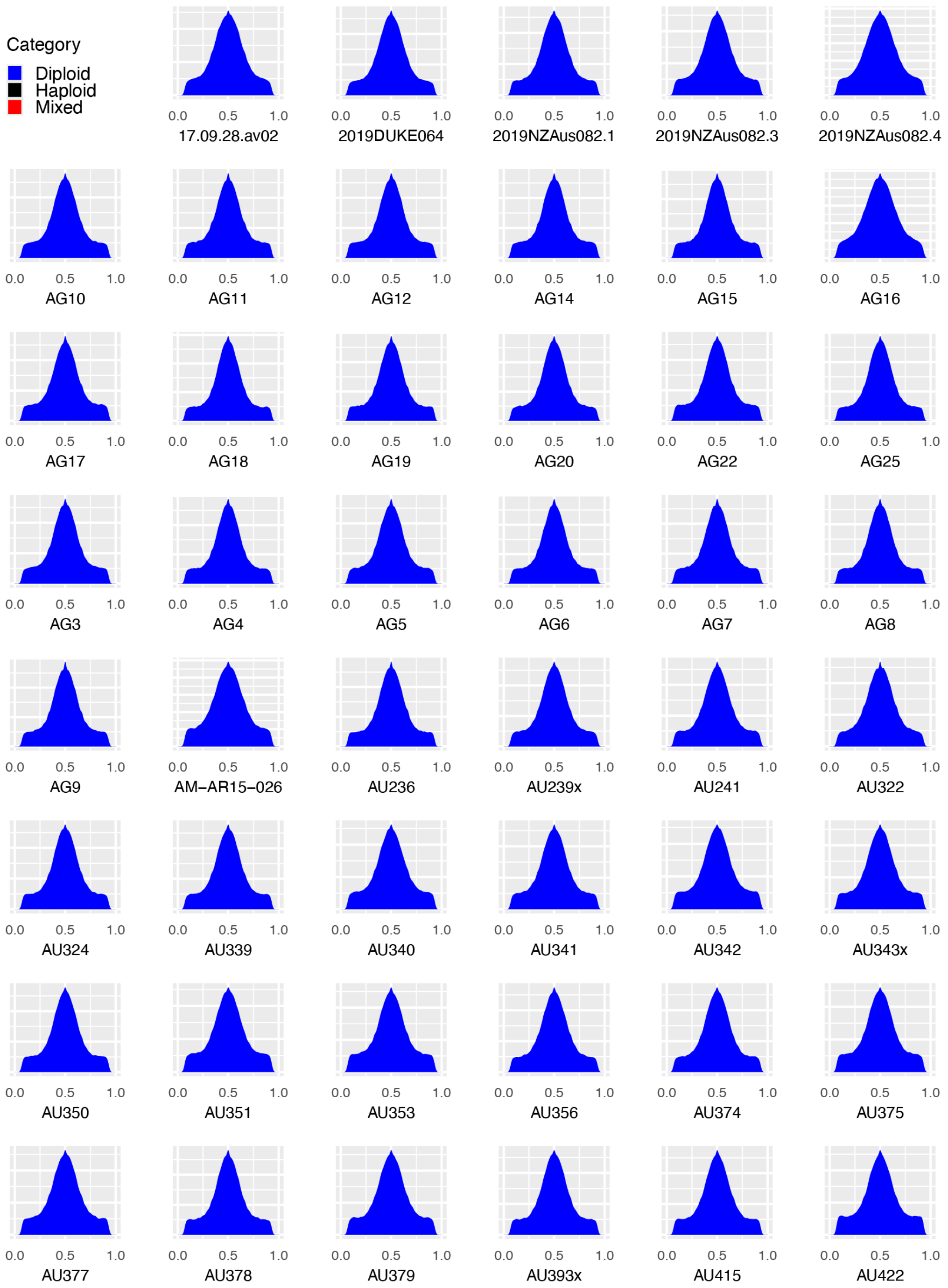


**Figure M2. Density of alternative allele frequency at heterozygous sites.**
One allele at each site is randomly picked as reference allele to make the density symmetric. Genomes labeled as haploid include the genomes whose heterozygous sites are below a hard cutoff 2% in Figure M1 and aurim734.

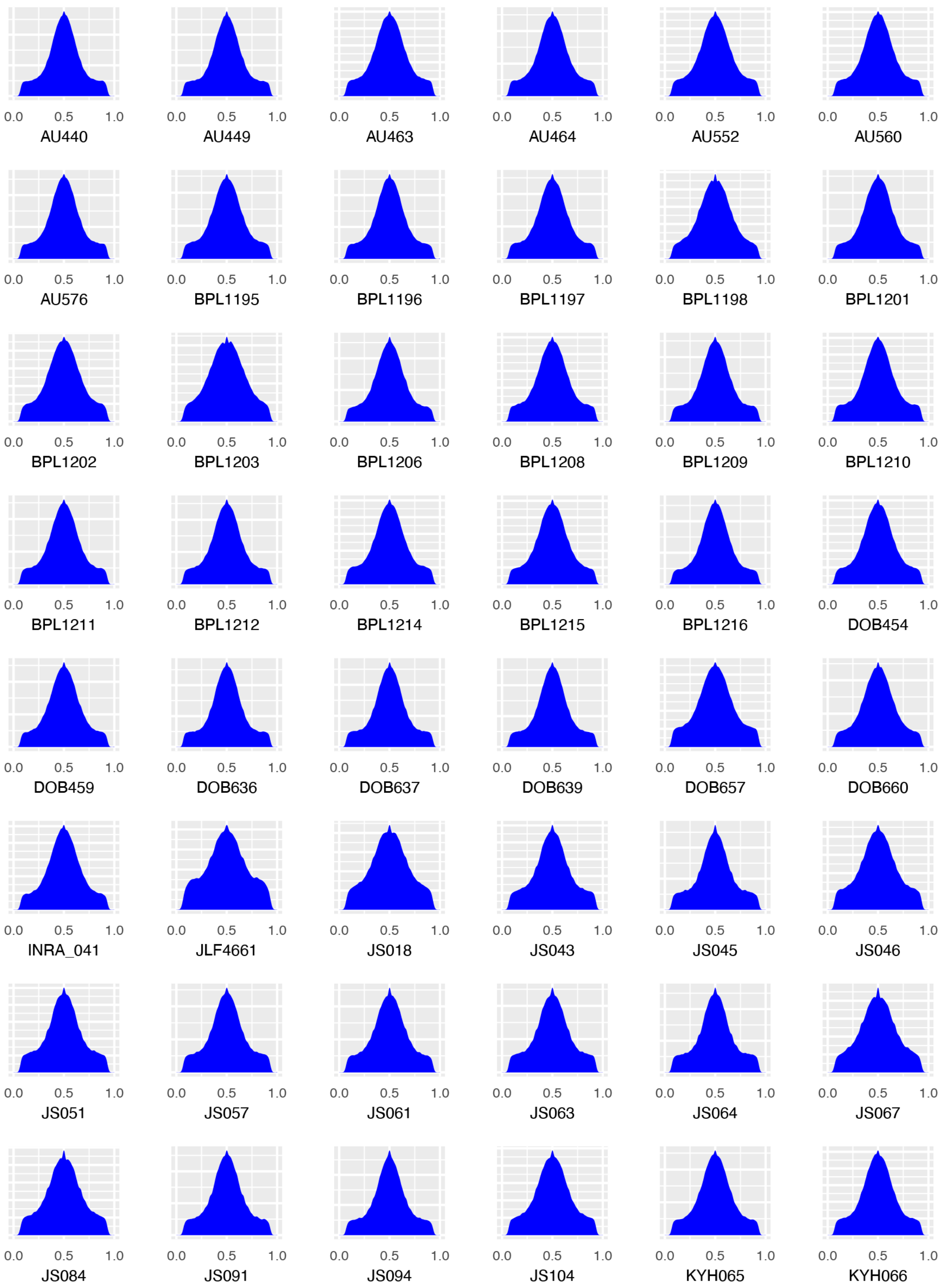


**Figure M2. Density of alternative allele frequency at heterozygous sites (continued).**

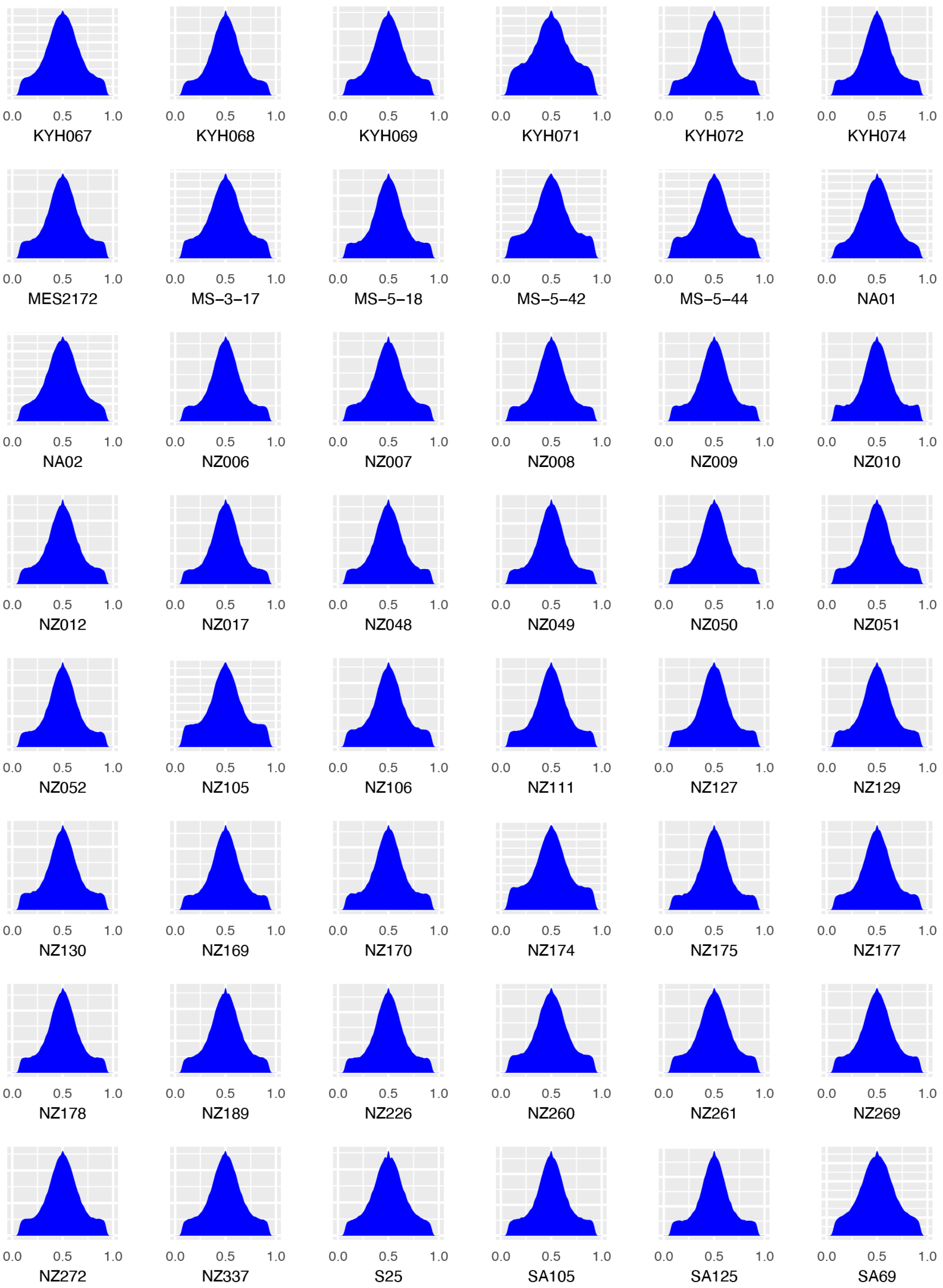


**Figure M2. Density of alternative allele frequency at heterozygous sites (continued).**

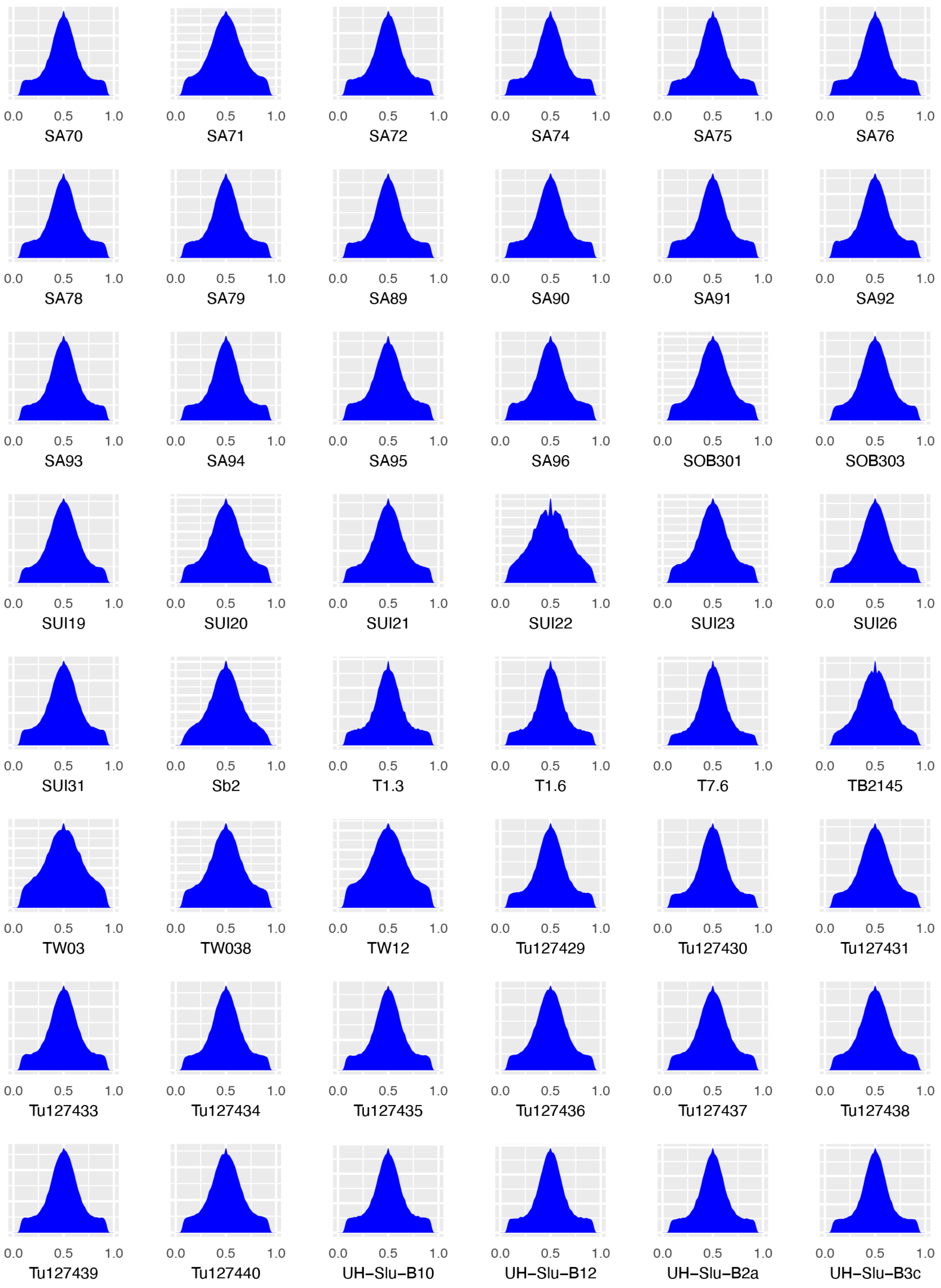


**Figure M2. Density of alternative allele frequency at heterozygous sites (continued).**

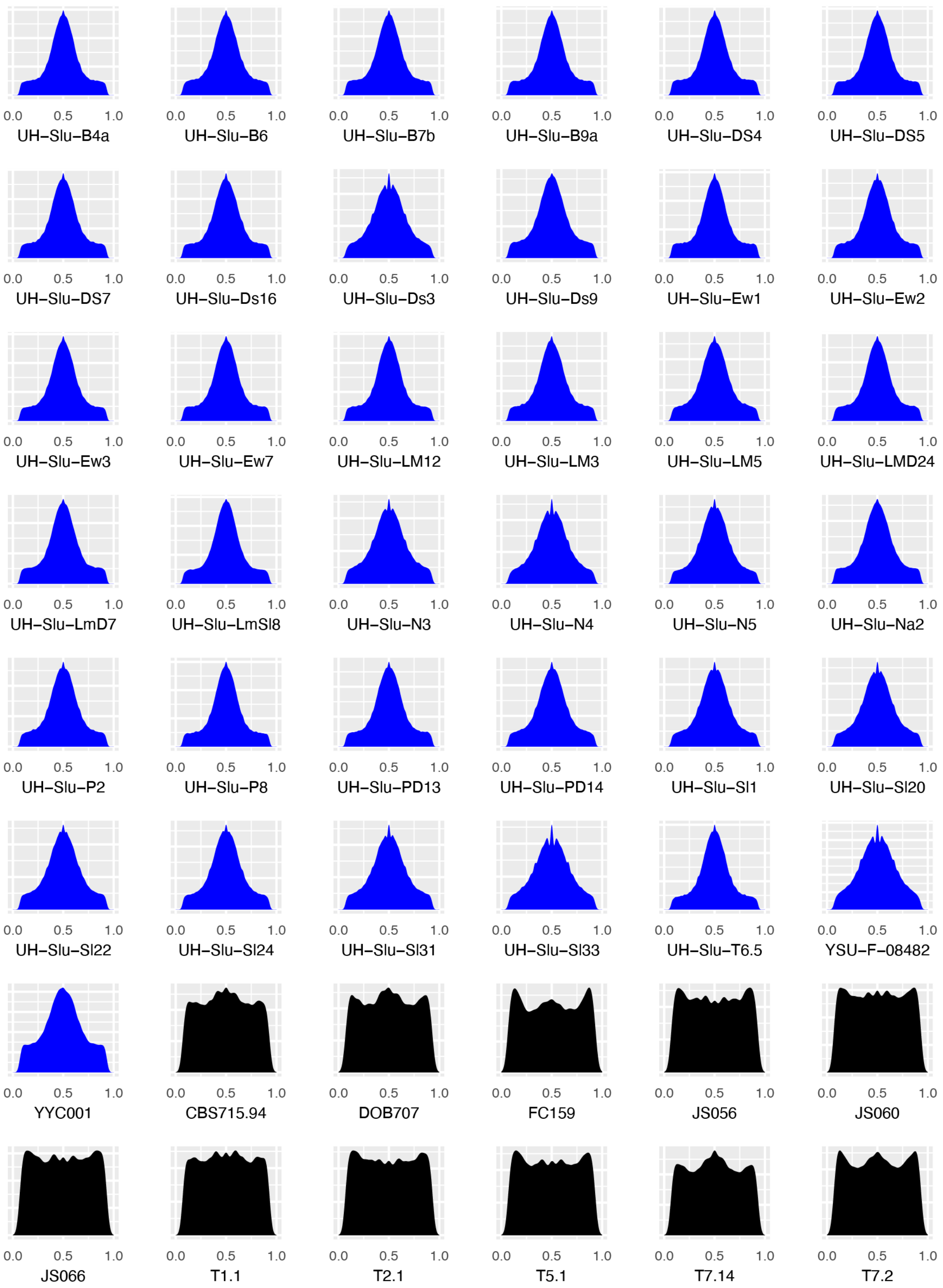


**Figure M2. Density of alternative allele frequency at heterozygous sites (continued).**

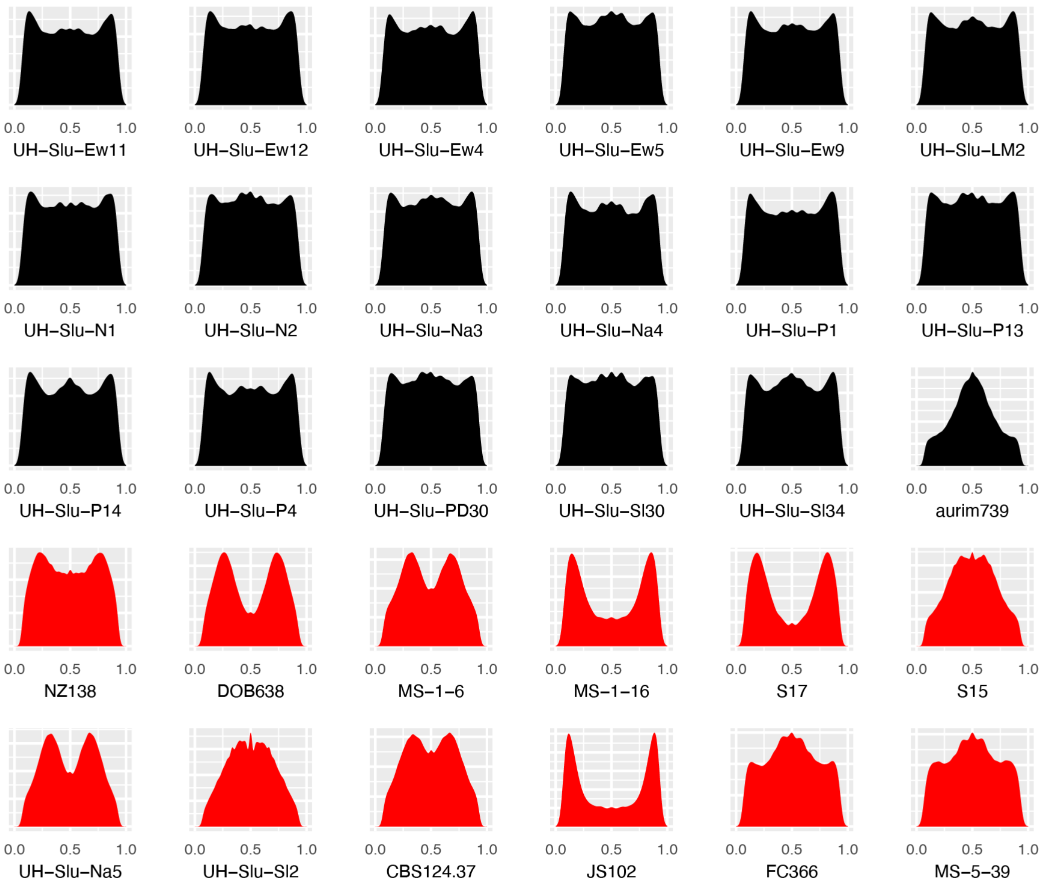


**Figure M2. Density of alternative allele frequency at heterozygous sites (continued).**

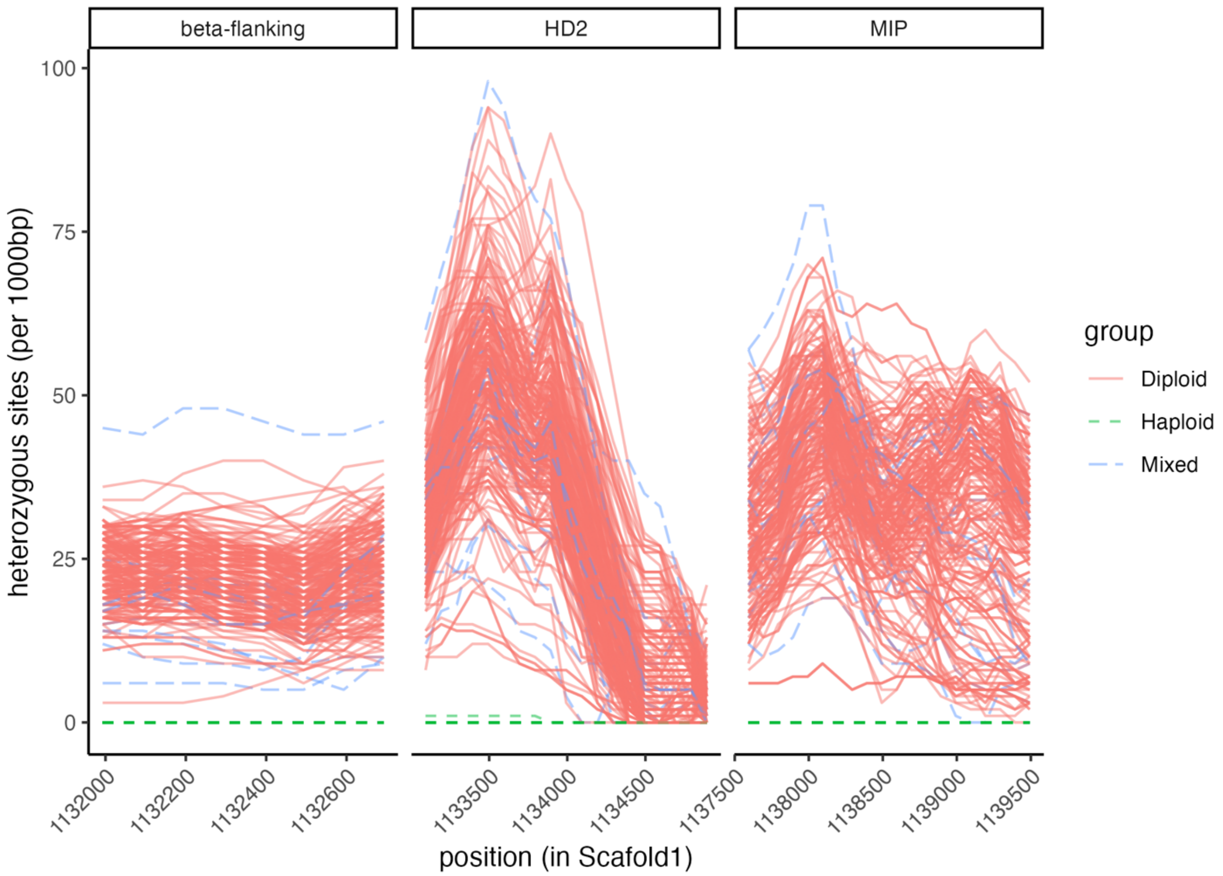


**Figure M3. Heterozygous site density around HD MAT mating type locus**

Each line represents the number of detected heterozygous sites of each genome in 1000bp windows around the mating type determining locus HD MAT locus. A diploid individual is expected to have two different mating type alleles at HD MAT locus, which creates heterozygous sites at HD MAT locus. Instead, a haploid individual is expected to have a single mating type, which does not present heterozygous sites. Genomes labeled as haploid include the genomes whose heterozygous sites are below a hard cutoff 2% in Figure M1 and aurim734.

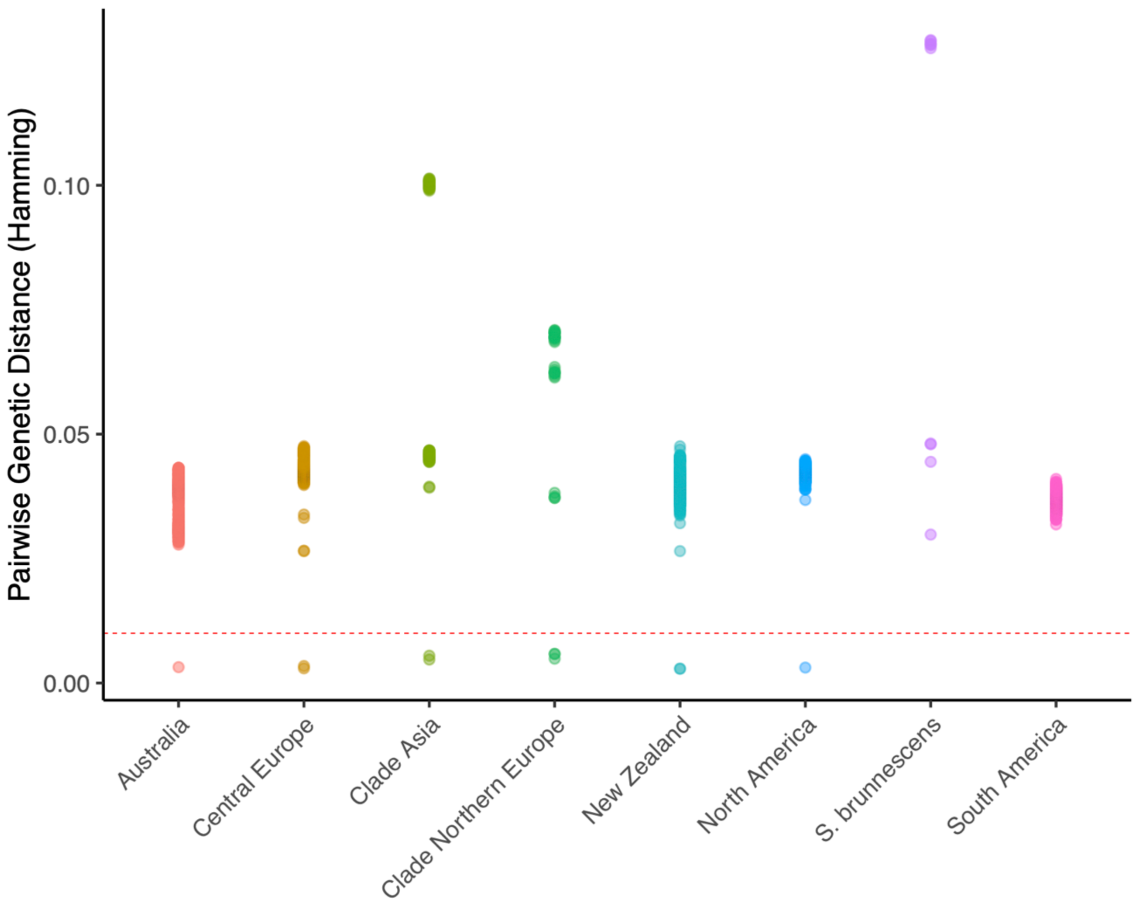


**Figure M4. Pairwise genetic distance between individuals in the same population.** Genetic distance is calculated from Hamming distance. Some genetic distance between genomes is nearly zero and are disjunct from other pairs in the same population, indicating they are likely to be the same genet. The hard cutoff is set to 0.01 as indicated by red dashed line. All the pairs of clonal genomes are not only collected in the same population but also the same locality.

**Invariability from sequencing methods and read length**

Although all the genomes included in the study were sequenced using the Illumina platform, two different read lengths, 150bp paired-end and 100bp paired-end, were applied through multiple batches. To verify the sequencing read lengths did not systematically affect genotyping, we bioinformatically assessed the differences in genotyping as if all the genomes had been sequenced in 100bp. The reads of all genomes sequenced in 150bp were trimmed to 100bp long by removing bases at the 3' end as if the sequencing did not extend further than 100bp. The same reference mapping and SNPs discovery methods were applied to the shortened reads as described in Method of the main text. The resultant genotypes from the shortened reads were then compared to the genotypes from the original reads.

The sequencing methods differing in read length among sequencing runs had marginal effects on the SNPs matrix. The differences in genotypes between original read length and 100bp reads were less than 1% among discovered SNPs for all genomes (Table M1). The discrepancy of genotypes in both alleles (i.e. changes from homozygous reference alleles to homozygous alternative alleles or vice versa) was less than 0.05%. The genotypes detected in the original read length but missed in 100bp reads were less than 2% of discovered SNPs. The principal components analysis was independently conducted to genotypes from original reads and shortened reads. The principal components from the two datasets showed clear collinearity (Figure M5), suggesting the differences in genotypes between the two types of reads were not systematic.

**Table M1. Discrepancy between genotypes detected from original reads (150bp) vs shortened reads (100bp) for genomes originally sequenced in 150bp.**

| genome ID | Number of SNPs called by both | Number of SNPs called by 150bp | Number of SNPs called by 100bp | Ratio of discrepant genotype (among sites called by both) | Ratio of homozygous discrepant genotype (among sites called by both) | Ratio of SNPs missed by 150bp | Ratio of SNPs missed by 100bp |
|---|---|---|---|---|---|---|---|
| 17.09.28.av02 | 720598 | 722424 | 721526 | 0.00773663 | 1.30E-04 | 0.00128616 | 0.0025276 |
| 2019DUKE064 | 723083 | 724132 | 724266 | 0.00747079 | 8.85E-05 | 0.00163338 | 0.00144863 |
| 2019NZAus082.1 | 722897 | 723789 | 724047 | 0.00719743 | 1.08E-04 | 0.00158829 | 0.0012324 |
| 2019NZAus082.3 | 720969 | 722066 | 722271 | 0.00746218 | 1.33E-04 | 0.00180265 | 0.00151925 |
| 2019NZAus082.4 | 723093 | 724739 | 724071 | 0.00805982 | 1.19E-04 | 0.0013507 | 0.00227116 |
| AG10 | 724147 | 724966 | 725131 | 0.00758133 | 8.98E-05 | 0.001357 | 0.00112971 |
| AG11 | 723374 | 724313 | 724490 | 0.00773735 | 9.40E-05 | 0.00154039 | 0.0012964 |
| AG12 | 724036 | 725046 | 725130 | 0.00718058 | 1.02E-04 | 0.00150869 | 0.00139302 |
| AG14 | 722844 | 723860 | 723902 | 0.00711633 | 1.01E-04 | 0.00146152 | 0.00140359 |
| AG15 | 724205 | 724936 | 725077 | 0.0074316 | 1.09E-04 | 0.00120263 | 0.00100836 |
| AG16 | 722003 | 723521 | 723419 | 0.00727836 | 8.31E-05 | 0.00195737 | 0.00209807 |
| AG17 | 722829 | 723841 | 723960 | 0.00730463 | 1.29E-04 | 0.00156224 | 0.0013981 |
| AG18 | 722165 | 723199 | 723167 | 0.00737228 | 1.02E-04 | 0.00138557 | 0.00142976 |
| AG19 | 722491 | 723432 | 723733 | 0.00743954 | 1.04E-04 | 0.0017161 | 0.00130074 |
| AG20 | 724323 | 725304 | 725361 | 0.00715427 | 7.59E-05 | 0.00143101 | 0.00135254 |
| AG22 | 723684 | 724595 | 724733 | 0.00760415 | 1.16E-04 | 0.00144743 | 0.00125725 |
| AG25 | 722861 | 723663 | 723909 | 0.00729186 | 1.18E-04 | 0.0014477 | 0.00110825 |
| AG3 | 723070 | 723887 | 724070 | 0.00738656 | 8.71E-05 | 0.00138108 | 0.00112863 |
| AG4 | 724138 | 724829 | 725068 | 0.00738809 | 9.39E-05 | 0.00128264 | 9.53E-04 |
| AG5 | 723077 | 724002 | 724111 | 0.00726479 | 9.82E-05 | 0.00142796 | 0.00127762 |
| AG6 | 722622 | 723499 | 723778 | 0.00699397 | 1.15E-04 | 0.00159717 | 0.00121216 |
| AG7 | 721845 | 722982 | 722985 | 0.00700843 | 1.25E-04 | 0.0015768 | 0.00157265 |
| AG8 | 723466 | 724305 | 724457 | 0.00757603 | 8.57E-05 | 0.00136792 | 0.00115835 |
| AG9 | 723704 | 724696 | 724830 | 0.0074326 | 1.05E-04 | 0.00155347 | 0.00136885 |
| AM-AR15-026 | 722030 | 723599 | 722974 | 0.00771021 | 1.09E-04 | 0.00130572 | 0.00216833 |
| AU236 | 722561 | 723604 | 723771 | 0.00738623 | 1.16E-04 | 0.0016718 | 0.0014414 |
| AU239x | 723629 | 724905 | 724637 | 0.00700912 | 1.31E-04 | 0.00139104 | 0.00176023 |
| AU241 | 724155 | 725142 | 725039 | 0.00726778 | 8.56E-05 | 0.00121924 | 0.00136111 |
| AU322 | 719807 | 721836 | 720946 | 0.00783127 | 1.42E-04 | 0.00157987 | 0.00281089 |
| AU324 | 723334 | 724299 | 724611 | 0.00727327 | 1.13E-04 | 0.00176232 | 0.00133232 |
| AU339 | 722802 | 723808 | 723874 | 0.00676672 | 8.16E-05 | 0.00148092 | 0.00138987 |
| AU340 | 721525 | 722889 | 722729 | 0.00698936 | 1.28E-04 | 0.00166591 | 0.00188687 |
| AU341 | 722174 | 723367 | 723258 | 0.00690277 | 1.05E-04 | 0.00149877 | 0.00164923 |

| | | | | | | | |
|---|---|---|---|---|---|---|---|
| AU342 | 721817 | 722945 | 722905 | 0.00662079 | 8.45E-05 | 0.00150504 | 0.00156028 |
| AU343x | 720979 | 722164 | 722531 | 0.00686705 | 1.35E-04 | 0.002148 | 0.0016409 |
| AU350 | 719649 | 720763 | 720831 | 0.00619747 | 1.06E-04 | 0.00163977 | 0.00154558 |
| AU351 | 722192 | 723687 | 723084 | 0.00683059 | 9.28E-05 | 0.0012336 | 0.00206581 |
| AU353 | 720817 | 722030 | 722380 | 0.00694074 | 1.17E-04 | 0.00216368 | 0.00167999 |
| AU356 | 722755 | 723691 | 723912 | 0.0067976 | 9.13E-05 | 0.00159826 | 0.00129337 |
| AU374 | 722569 | 723750 | 723681 | 0.0063385 | 1.07E-04 | 0.00153659 | 0.00163178 |
| AU375 | 723713 | 724672 | 724711 | 0.00664075 | 9.53E-05 | 0.0013771 | 0.00132336 |
| AU377 | 723472 | 724449 | 724511 | 0.00686689 | 9.68E-05 | 0.00143407 | 0.00134861 |
| AU378 | 721945 | 723067 | 723368 | 0.00677198 | 9.70E-05 | 0.00196719 | 0.00155172 |
| AU379 | 721741 | 722806 | 722961 | 0.0069083 | 9.56E-05 | 0.0016875 | 0.00147342 |
| AU393x | 721431 | 722686 | 722583 | 0.00658414 | 7.90E-05 | 0.00159428 | 0.00173658 |
| AU415 | 720335 | 721747 | 721511 | 0.00671493 | 8.88E-05 | 0.00162991 | 0.00195636 |
| AU422 | 724445 | 725390 | 725554 | 0.00744156 | 9.80E-05 | 0.00152849 | 0.00130275 |
| AU440 | 723219 | 724121 | 724179 | 0.00713753 | 7.60E-05 | 0.00132564 | 0.00124565 |
| AU449 | 722030 | 723067 | 723234 | 0.00733903 | 9.42E-05 | 0.00166474 | 0.00143417 |
| AU463 | 722314 | 723436 | 723414 | 0.0078027 | 1.38E-04 | 0.00152057 | 0.00155093 |
| AU552 | 722428 | 723710 | 723428 | 0.00784576 | 1.16E-04 | 0.00138231 | 0.00177143 |
| AU560 | 722394 | 723939 | 723434 | 0.00751529 | 1.12E-04 | 0.00143759 | 0.00213416 |
| AU576 | 723124 | 724235 | 724358 | 0.00709284 | 9.82E-05 | 0.00170358 | 0.00153403 |
| BPL1195 | 722984 | 723961 | 724095 | 0.00750224 | 1.05E-04 | 0.00153433 | 0.00134952 |
| BPL1196 | 723198 | 724243 | 724329 | 0.00763138 | 1.23E-04 | 0.00156145 | 0.00144289 |
| BPL1197 | 721852 | 722859 | 723172 | 0.00721893 | 1.21E-04 | 0.00182529 | 0.00139308 |
| BPL1198 | 720940 | 723162 | 722092 | 0.00894943 | 1.19E-04 | 0.00159536 | 0.00307262 |
| BPL1201 | 721947 | 723125 | 723327 | 0.00774295 | 1.27E-04 | 0.00190785 | 0.00162904 |
| BPL1202 | 722353 | 723282 | 723827 | 0.0079864 | 9.55E-05 | 0.0020364 | 0.00128442 |
| BPL1203 | 721244 | 723426 | 722571 | 0.00885831 | 1.30E-04 | 0.0018365 | 0.0030162 |
| BPL1206 | 723088 | 724429 | 724004 | 0.00767403 | 1.33E-04 | 0.00126519 | 0.00185111 |
| BPL1208 | 724020 | 725379 | 724901 | 0.00822767 | 1.22E-04 | 0.00121534 | 0.0018735 |
| BPL1209 | 723709 | 724596 | 724846 | 0.0077697 | 9.81E-05 | 0.00156861 | 0.00122413 |
| BPL1210 | 721200 | 722499 | 722549 | 0.00741681 | 1.03E-04 | 0.001867 | 0.00179793 |
| BPL1211 | 723086 | 723915 | 724170 | 0.0079382 | 1.01E-04 | 0.00149689 | 0.00114516 |
| BPL1212 | 721691 | 722942 | 722821 | 0.0073785 | 0.00012609 | 0.00156332 | 0.00173043 |
| BPL1214 | 720603 | 722422 | 721747 | 0.00796278 | 1.07E-04 | 0.00158504 | 0.00251792 |
| BPL1215 | 723093 | 724558 | 724080 | 0.00824514 | 9.96E-05 | 0.00136311 | 0.00202192 |
| BPL1216 | 722151 | 723230 | 723273 | 0.00790001 | 9.28E-05 | 0.00155128 | 0.00149192 |
| DOB454 | 720385 | 721951 | 721616 | 0.00812205 | 1.22E-04 | 0.00170589 | 0.00216912 |
| DOB459 | 723264 | 724153 | 724457 | 0.00761686 | 1.02E-04 | 0.00164675 | 0.00122764 |
| DOB636 | 724573 | 725419 | 725479 | 0.00767348 | 9.52E-05 | 0.00124883 | 0.00116622 |

| | | | | | | | |
|---|---|---|---|---|---|---|---|
| DOB639 | 723376 | 724247 | 724346 | 0.00729911 | 8.99E-05 | 0.00133914 | 0.00120263 |
| DOB657 | 722583 | 724109 | 723511 | 0.00867444 | 9.83E-05 | 0.00128263 | 0.00210742 |
| DOB660 | 724790 | 725587 | 725932 | 0.00804233 | 1.15E-04 | 0.00157315 | 0.00109842 |
| INRA_041 | 724012 | 724960 | 725095 | 0.0077844 | 1.04E-04 | 0.0014936 | 0.00130766 |
| JS018 | 688796 | 696005 | 693909 | 0.00861648 | 3.48E-04 | 0.0073684 | 0.01035768 |
| JS043 | 688342 | 693899 | 693483 | 0.00821249 | 3.07E-04 | 0.0074133 | 0.00800837 |
| JS045 | 689794 | 694523 | 695274 | 0.00748919 | 2.46E-04 | 0.00788178 | 0.00680899 |
| JS046 | 689612 | 695509 | 694388 | 0.0080451 | 3.15E-04 | 0.006878 | 0.00847868 |
| JS051 | 689570 | 694232 | 694907 | 0.0076758 | 2.97E-04 | 0.00768016 | 0.00671533 |
| JS057 | 686036 | 692373 | 690934 | 0.007555 | 2.80E-04 | 0.00708895 | 0.00915258 |
| JS061 | 685660 | 692694 | 690300 | 0.00805939 | 3.50E-04 | 0.00672172 | 0.01015456 |
| JS063 | 687712 | 693158 | 692939 | 0.00749296 | 3.11E-04 | 0.00754323 | 0.00785679 |
| JS067 | 682980 | 691091 | 687998 | 0.00817886 | 3.63E-04 | 0.00729363 | 0.01173652 |
| JS084 | 686054 | 693331 | 690747 | 0.00814805 | 3.08E-04 | 0.00679409 | 0.01049571 |
| KYH065 | 723639 | 724454 | 724706 | 0.00791279 | 8.84E-05 | 0.00147232 | 0.00112499 |
| KYH066 | 721893 | 722921 | 723138 | 0.00783496 | 1.05E-04 | 0.00172166 | 0.00142201 |
| KYH067 | 722156 | 723601 | 723279 | 0.00806059 | 1.08E-04 | 0.00155265 | 0.00199696 |
| KYH068 | 723527 | 724471 | 724748 | 0.00752702 | 1.02E-04 | 0.00168472 | 0.00130302 |
| KYH069 | 723194 | 724575 | 724259 | 0.00786511 | 1.16E-04 | 0.00147047 | 0.00190594 |
| KYH072 | 721476 | 722466 | 722698 | 0.00729893 | 9.84E-05 | 0.00169089 | 0.00137031 |
| KYH074 | 720787 | 721757 | 722125 | 0.00701455 | 9.43E-05 | 0.00185286 | 0.00134394 |
| MES2172 | 722950 | 724001 | 724079 | 0.00795214 | 1.15E-04 | 0.00155922 | 0.00145166 |
| NA01 | 721145 | 722400 | 722381 | 0.00683358 | 1.10E-04 | 0.00171101 | 0.00173726 |
| NA02 | 719612 | 721141 | 720847 | 0.00653964 | 9.45E-05 | 0.00171326 | 0.00212025 |
| NZ006 | 721834 | 722918 | 722884 | 0.00718725 | 1.03E-04 | 0.00145252 | 0.00149948 |
| NZ007 | 724263 | 725074 | 725386 | 0.00737992 | 1.01E-04 | 0.00154814 | 0.00111851 |
| NZ009 | 723810 | 724820 | 724934 | 0.00787776 | 9.39E-05 | 0.00155049 | 0.00139345 |
| NZ010 | 723851 | 725339 | 724666 | 0.00846307 | 9.81E-05 | 0.00112466 | 0.00205145 |
| NZ012 | 722497 | 723655 | 723815 | 0.00789484 | 1.18E-04 | 0.00182091 | 0.00160021 |
| NZ017 | 721856 | 723414 | 722965 | 0.00701248 | 9.70E-05 | 0.00153396 | 0.00215368 |
| NZ048 | 722853 | 723825 | 723962 | 0.00757415 | 1.29E-04 | 0.00153185 | 0.00134287 |
| NZ049 | 722764 | 723747 | 723815 | 0.00688053 | 1.15E-04 | 0.00145203 | 0.00135821 |
| NZ050 | 724380 | 725155 | 725458 | 0.00734007 | 9.11E-05 | 0.00148596 | 0.00106874 |
| NZ051 | 723771 | 724965 | 724802 | 0.00749685 | 8.98E-05 | 0.00142246 | 0.00164698 |
| NZ052 | 723045 | 724028 | 724341 | 0.0069885 | 9.82E-05 | 0.00178921 | 0.00135768 |
| NZ105 | 718075 | 719764 | 719531 | 0.00595202 | 1.17E-04 | 0.00202354 | 0.0023466 |
| NZ106 | 720692 | 721792 | 722081 | 0.00669079 | 1.30E-04 | 0.00192361 | 0.00152398 |
| NZ111 | 722012 | 723287 | 723082 | 0.00733506 | 1.08E-04 | 0.00147978 | 0.00176279 |
| NZ127 | 722837 | 723789 | 723868 | 0.0070832 | 1.13E-04 | 0.00142429 | 0.0013153 |

| | | | | | | | |
|---|---|---|---|---|---|---|---|
| NZ129 | 721541 | 722492 | 722774 | 0.00681735 | 9.42E-05 | 0.00170593 | 0.00131628 |
| NZ130 | 724253 | 725187 | 725231 | 0.00743249 | 8.70E-05 | 0.00134854 | 0.00128794 |
| NZ169 | 722708 | 723820 | 723714 | 0.0072173 | 9.96E-05 | 0.00139005 | 0.00153629 |
| NZ170 | 722130 | 723203 | 723304 | 0.00745157 | 9.69E-05 | 0.00162311 | 0.00148368 |
| NZ174 | 717897 | 720016 | 719253 | 0.00622513 | 1.17E-04 | 0.00188529 | 0.00294299 |
| NZ177 | 722070 | 723374 | 723135 | 0.0076073 | 1.27E-04 | 0.00147275 | 0.00180266 |
| NZ178 | 722568 | 723928 | 723656 | 0.00719932 | 1.22E-04 | 0.00150348 | 0.00187864 |
| NZ189 | 723981 | 725274 | 724872 | 0.00788418 | 9.12E-05 | 0.00122918 | 0.00178277 |
| NZ226 | 722242 | 723374 | 723275 | 0.00708904 | 9.97E-05 | 0.00142823 | 0.00156489 |
| NZ260 | 720676 | 722017 | 721756 | 0.00734588 | 1.18E-04 | 0.00149635 | 0.0018573 |
| NZ261 | 721618 | 723203 | 722769 | 0.00802086 | 1.46E-04 | 0.00159249 | 0.00219164 |
| NZ269 | 723026 | 724009 | 724251 | 0.0071021 | 9.13E-05 | 0.0016914 | 0.00135772 |
| NZ272 | 721633 | 722856 | 722969 | 0.00731674 | 1.25E-04 | 0.00184794 | 0.0016919 |
| NZ337 | 722936 | 723909 | 723997 | 0.00779875 | 1.16E-04 | 0.00146548 | 0.00134409 |
| S25 | 722502 | 724398 | 723438 | 0.00847472 | 1.44E-04 | 0.00129382 | 0.00261735 |
| SA105 | 723287 | 724266 | 724541 | 0.00739679 | 1.05E-04 | 0.00173075 | 0.00135171 |
| SA125 | 723563 | 724376 | 724525 | 0.00722397 | 7.74E-05 | 0.00132777 | 0.00112235 |
| SA69 | 722754 | 724046 | 723857 | 0.00666894 | 8.02E-05 | 0.00152378 | 0.00178442 |
| SA70 | 722651 | 723576 | 723722 | 0.00779906 | 7.75E-05 | 0.00147985 | 0.00127837 |
| SA71 | 722462 | 723601 | 723774 | 0.00667025 | 8.44E-05 | 0.00181272 | 0.00157407 |
| SA72 | 722797 | 723785 | 723968 | 0.00716937 | 9.27E-05 | 0.00161747 | 0.00136505 |
| SA74 | 723013 | 723828 | 724011 | 0.00742587 | 8.58E-05 | 0.00137843 | 0.00112596 |
| SA75 | 722102 | 723109 | 723233 | 0.00740616 | 9.97E-05 | 0.00156381 | 0.0013926 |
| SA76 | 724059 | 725001 | 724977 | 0.00778666 | 1.12E-04 | 0.00126625 | 0.00129931 |
| SA78 | 722812 | 723833 | 723927 | 0.00777104 | 9.27E-05 | 0.00154021 | 0.00141055 |
| SA79 | 723447 | 724242 | 724473 | 0.00726107 | 8.85E-05 | 0.0014162 | 0.0010977 |
| SA89 | 723527 | 724328 | 724454 | 0.00742889 | 7.46E-05 | 0.00127958 | 0.00110585 |
| SA90 | 723127 | 724112 | 724251 | 0.00753533 | 1.12E-04 | 0.00155195 | 0.00136029 |
| SA91 | 723597 | 724549 | 724761 | 0.00760506 | 9.81E-05 | 0.00160605 | 0.00131392 |
| SA92 | 722239 | 723275 | 723485 | 0.00719291 | 1.08E-04 | 0.00172222 | 0.00143237 |
| SA93 | 722572 | 723491 | 723655 | 0.00758956 | 1.26E-04 | 0.00149657 | 0.00127023 |
| SA94 | 723366 | 724180 | 724326 | 0.00746372 | 1.05E-04 | 0.00132537 | 0.00112403 |
| SA95 | 723763 | 724724 | 725075 | 0.00770694 | 1.22E-04 | 0.00180947 | 0.00132602 |
| SA96 | 723262 | 724115 | 724464 | 0.0078616 | 7.47E-05 | 0.00165916 | 0.00117799 |
| SOB301 | 722696 | 723931 | 723967 | 0.00735579 | 1.31E-04 | 0.0017556 | 0.00170596 |
| SOB303 | 721142 | 722214 | 722301 | 0.00771831 | 1.08E-04 | 0.00160459 | 0.00148432 |
| SUI19 | 722555 | 723465 | 723646 | 0.00743888 | 1.13E-04 | 0.00150764 | 0.00125784 |
| SUI20 | 690991 | 695922 | 695035 | 0.00900012 | 2.74E-04 | 0.00581841 | 0.00708556 |
| SUI21 | 691717 | 695650 | 696275 | 0.00866106 | 2.05E-04 | 0.00654626 | 0.00565371 |

| | | | | | | | |
|---|---|---|---|---|---|---|---|
| SUI26 | 722292 | 723357 | 723599 | 0.00758834 | 1.22E-04 | 0.00180625 | 0.0014723 |
| SUI31 | 722665 | 723723 | 723868 | 0.00801063 | 1.26E-04 | 0.00166191 | 0.00146189 |
| T1.3 | 694088 | 698088 | 698469 | 0.00911988 | 2.43E-04 | 0.00627229 | 0.00572994 |
| T1.6 | 693707 | 697590 | 698254 | 0.00851656 | 2.31E-04 | 0.00651196 | 0.00556631 |
| T7.6 | 696653 | 700192 | 700838 | 0.00857098 | 2.24E-04 | 0.00597142 | 0.00505433 |
| TB2145 | 690912 | 697156 | 694780 | 0.00928917 | 2.68E-04 | 0.00556723 | 0.00895639 |
| TW03 | 680317 | 686421 | 685416 | 0.00733482 | 2.78E-04 | 0.00743928 | 0.0088925 |
| TW038 | 679645 | 685440 | 685118 | 0.00786881 | 2.96E-04 | 0.0079884 | 0.00845442 |
| Tu127429 | 723395 | 724351 | 724361 | 0.00738186 | 8.29E-05 | 0.00133359 | 0.0013198 |
| Tu127430 | 720275 | 721633 | 721411 | 0.00734025 | 9.30E-05 | 0.00157469 | 0.00188184 |
| Tu127431 | 720768 | 722393 | 721898 | 0.00766127 | 1.32E-04 | 0.00156532 | 0.00224947 |
| Tu127433 | 722683 | 723663 | 723676 | 0.0075787 | 9.55E-05 | 0.00137216 | 0.00135422 |
| Tu127434 | 720587 | 721534 | 721807 | 0.00727879 | 1.14E-04 | 0.0016902 | 0.00131248 |
| Tu127435 | 721623 | 722577 | 722595 | 0.00727111 | 8.73E-05 | 0.00134515 | 0.00132027 |
| Tu127436 | 722046 | 723556 | 723058 | 0.00762832 | 1.05E-04 | 0.00139961 | 0.00208692 |
| Tu127437 | 721373 | 722676 | 722506 | 0.00758553 | 1.16E-04 | 0.00156815 | 0.00180302 |
| Tu127438 | 721417 | 722900 | 722481 | 0.00778745 | 1.05E-04 | 0.0014727 | 0.00205146 |
| Tu127439 | 723840 | 724592 | 724775 | 0.00717562 | 1.11E-04 | 0.00129006 | 0.00103783 |
| Tu127440 | 721590 | 723270 | 722598 | 0.00818055 | 1.22E-04 | 0.00139497 | 0.00232278 |
| YSU-F-08482 | 718497 | 721710 | 719850 | 0.01069176 | 1.60E-04 | 0.00187956 | 0.00445193 |
| YYC001 | 713336 | 714877 | 715127 | 0.006471 | 1.40E-04 | 0.00250445 | 0.00215562 |

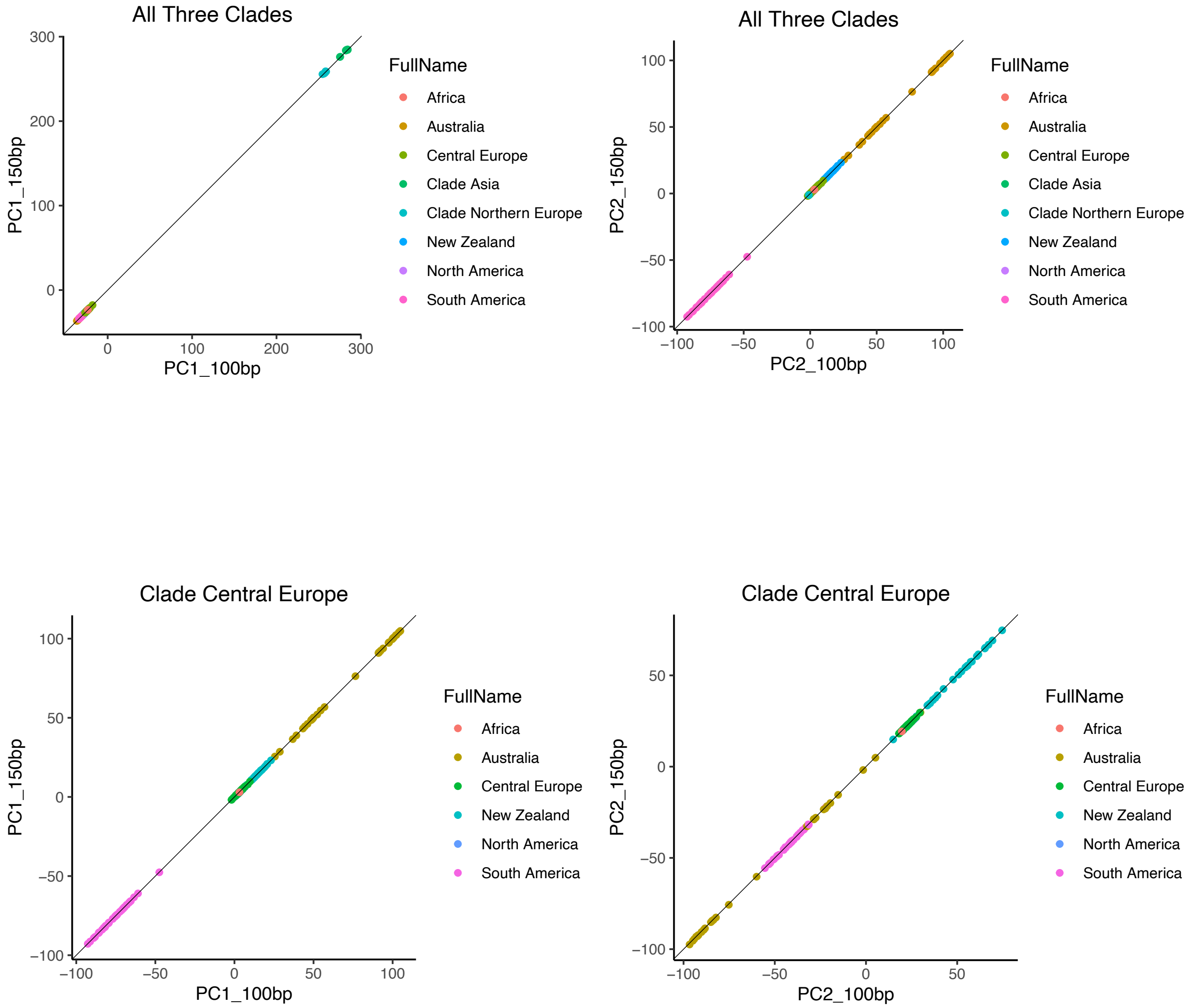


**Figure M5. Comparison of principal component values between genotypes detected from original reads (150bp) vs shortened reads (100bp).**

Each point represents a single genome sequenced in 150bp. The x-axis and y-axis show their respective principal component values based on genotypes detected from original reads (150bp) and shortened reads (100bp). Strong collinearity suggests no systematic differences between genotypes from different sequencing methods.

#### Invariability of PCA results from missing SNP data

Since the PCA calculation in dudi.pca replaces missing data with the mean of the variable, potentially distorting the calculation in unpredictable ways, we investigated the PCA without missing data. By selecting 315,052 SNPs that are present across all genomes for an alternative PCA, the first two principal components from the two datasets showed clear collinearity (Figure M6), confirming that the treatment of missing data does not systematically change the results.

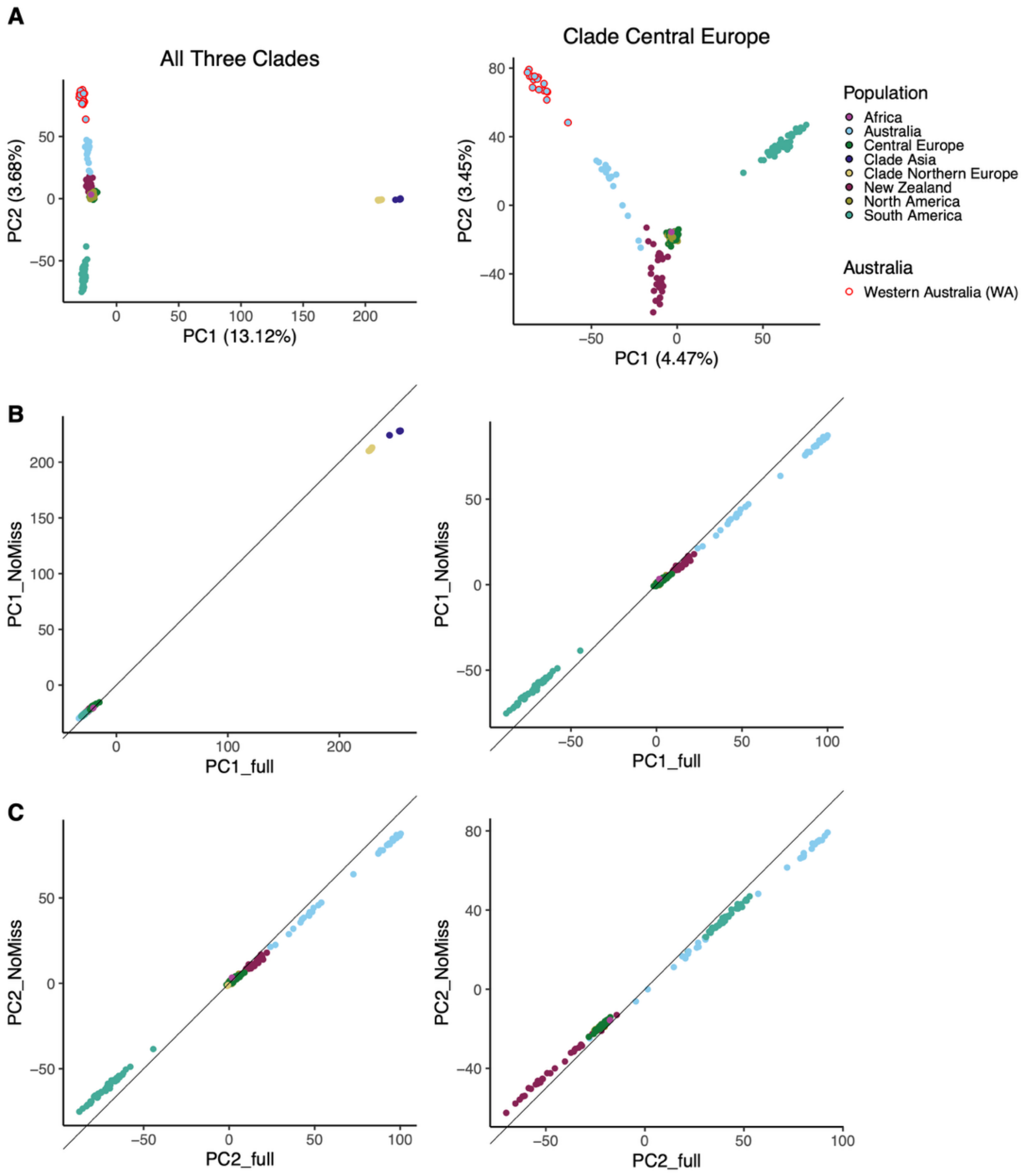


**Figure M6. Comparison of principal component values between datasets with or without missing genotypes.**

Each point represents the PC values of a single genome. The x-axis and y-axis show their respective principal component values based on genotypes of all SNPs and SNPs without missing values. Strong collinearity suggests no systematic differences between datasets with or without missing SNPs.

**Comparison of different window sizes of Tajima's D calculation**

We explored the impact of Tajima's D calculated by various arbitrary window sizes. The variation of Tajima's D across the genome depends on the choice of window sizes, where smaller window sizes lead to higher variation and alter the significance level because of it (Figure M7). However, the choice of window sizes does not change the sign and median of Tajima's D calculation.

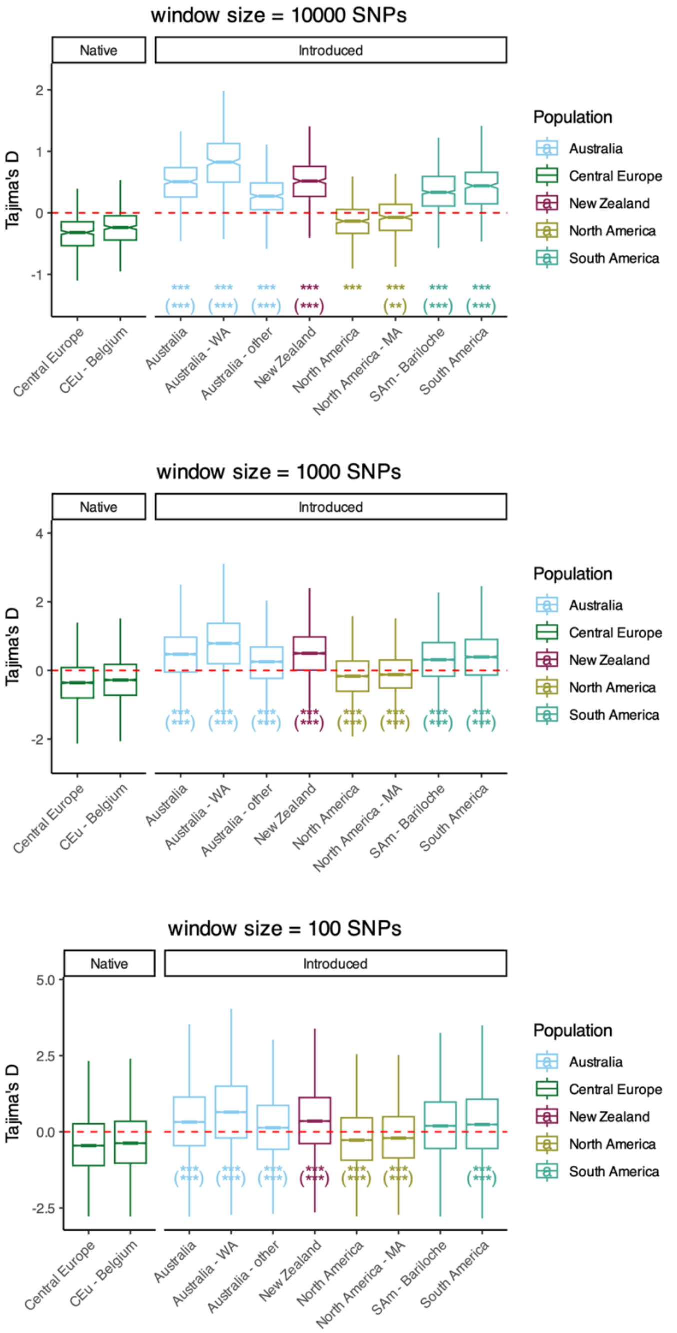


**Figure M7. Comparison of different window sizes of Tajima's D calculation.** The same calculation as in Figure 5 but use various window sizes. The significance of each introduced population compared to Central Europe was labeled as $*$ for $p < 0.05$, $**$ for $p < 0.01$, and $***$ for $p < 0.001$. The comparison to the subpopulation of Central Europe in Belgium (CEu-Belgium) was shown inside parenthesis.

**Extensive methods of demographic modeling**

Demographic modeling with momi2 (Kamm *et al.*, 2020) was used to explicitly infer the divergence history and growth rate parameters based on the site frequency spectrum (SFS). Momi2 applies composite likelihood inference for models including divergence, population size, migration, and growth rate for multiple populations at the same time. The mutation rate was set to $10^{-7}$ $year^{-1}$ $site^{-1}$ as used in another introduced basidiomycete fungus, *Serpula lacrymans* (Skrede *et al.*, 2021). The generation time was assumed to be 10 years according to the fact that most *Suillus luteus* reproduce at trees 5-15 years old (Shaw *et al.*, 2003). The clades beyond the introduction scene (Clade Asia and Clade Northern Europe, see Results) were treated as outgroup to calculate unfolded SFS for Clade Central Europe using functions *read_vcf* and *extract_sfs* in *momi2.* To balance the sample size among populations, the largest subpopulation in each population was treated as a representative deme for inference when possible, namely Belgium in Europe, Bariloche in South America, and Massachusetts in North America. The method used for optimizing likelihoods of all models in this research was truncated newton method (tnc) as recommended in the user's manual of *momi2*.

The population size of Europe ($N_{Eu}$) was estimated by implementing a model containing a single population (A0). The optimized $N_{Eu}$ converged to 3339.5 from a wide range of initial states from $N_{Eu}$=30 to $N_{Eu}$=300000 (Figure M8). A simple model was applied to estimate the divergence time and population growth rate (Figure S1). Inference was applied for each introduced population at one time. The model (A1) had a single divergence event from Europe to introduced population without growth, which contained four parameters: population size of Europe ($N_{Eu}$), population size of introduced population ($N_{in}$), time of divergence from Europe to introduced population ($t_{in}$), constant growth rate per year of introduced population since the time of divergence to date ($r_{in}$). The initial states of parameters were $N_{Eu}$=3,000, $N_{in}$=500, $t_{in}$=200, and $g_{in}$=0 for this model. The lower and upper boundaries of $r_{in}$ were set to 0.0 and 0.1, respectively. The fold of growth (ratio of current population size to the initial population size of introduction) was calculated by the exponential growth since the time of introduction as $(1 + r_{in})^{t_{in}}$. The initial state of $N_{Eu}$ was based on the approximate convergence value of $N_{Eu}$ at 3,000 in model A0, $t_{in}$ was based on the known time of introduction, and $N_{in}$ was based on the relative moderate decrease in population size according to observed nucleotide diversity (Figure 5 in the main text). Alternative values of the initial states of $t_{in}$ and $N_{in}$ were explored, but they had little effect on the inferences (Figure M9).

To test the possibility of ancient dispersal before human activity, an alternative model (A1a) with the same construction, except for a hard constraint of divergence time more than 1000 generations was compared to exclude the possibility of long divergence from Europe to an introduced population that predates the introduction of pines. The initial states of parameters were $N_{Eu}$=3,000, $N_{in}$=500, and $g_{in}$=0 as in the A1 model.

Additional models (B1-B3) were used to test the hypotheses of sequential divergence of introduced populations. Those models contained the population in Europe and two introduced populations at once to test whether the sequential divergence scenarios better explain the divergence history.

The first model (B1) was designed to fit the scenario of independent introduction. The model designated two introduced populations that independently diverged from the Europe population some time ago. This model contained two divergence events with 7 parameters, including population size of Europe ($N_{Eu}$), population size of the first introduced population ($N_{in1}$), time of divergence from Europe population to first introduced population ($t_{in1}$), constant growth rate per year of the first introduced population since the time of divergence to date ($r_{in1}$), population size of the second introduced population ($N_{in2}$), time of divergence from Europe population to second introduced population ($t_{in2}$), and constant population growth rate per year of the second introduced population since the time of divergence to date ($r_{in2}$). The same initial states of Europe and the introduced populations were set as in model A1 ($N_{Eu}$=3,000, $N_{in1}$=500, $t_{in1}$=200, $r_{in1}$=0, $N_{in2}$=500, $t_{in2}$=200, $r_{in2}$=0). The lower and upper boundaries of $r_{in1}$ and $r_{in2}$ were set to 0.0 and 0.1, respectively, as in model A1.

The second model (B2) was designed to fit the scenario of sequential introduction. It designated one introduced population diverged from the Europe population followed by the other population diverged from the first introduced population. The same parameters were contained in model B2 as in model B1, but the topology of divergences differed from B1 (Figure S1). The initial states of model B2 were the same as B1, except that the divergence time of the second population was set to half of the preliminarily optimized divergence time of the first population. $t_{in2}$ was constrained lower than $t_{in1}$ as implied by the topology.

The third model (B3) was designed to fit the scenario of independent introduction with later admixture. It designates two introduced populations independently diverged from the Europe population as model B1, followed by a single pulse of unidirectional migration from one introduced population to the other introduced population. In total 9 parameters were contained in Model B3, including the same 7 parameters in Model B1 and Model B2, plus time of migration ($t_m$) and portion of migration (m). The initial states of the two additional parameters were m=0.01 and $t_m$= $t_{in2}$/2. To distinguish migration from divergence events, the migration was limited to the minority of the receiving population, i.e., the upper boundary of m was set to 0.5. $t_m$ was constrained lower than $t_{in1}$ and $t_{in2}$ as implied by the topology. Other parameters had the same initial states as in models B1 and B2.

Sequential optimization was performed in model B1-B3. The divergence event of the first introduced population was optimized before adding the divergence event of the second introduced population, without population growth rates in both introduced populations. The migration event was then added and optimized when applicable. Lastly, the population growth rates in both introduced populations were added to the optimized model for the final optimization.

We investigated the models and confirmed that the mutation rate does not change the likelihood calculation. In the diffusion-based modeling of momi2, the mutation rate is solely used to determine the scaling of population size and divergence time. When the mutation rate is increased or decreased 10 times, the likelihoods are exactly the same when the population size and divergence time are scaled by 10 times accordingly. The calculated likelihood of such scaling for the best models of the investigated populations is shown in Table M2 as examples.

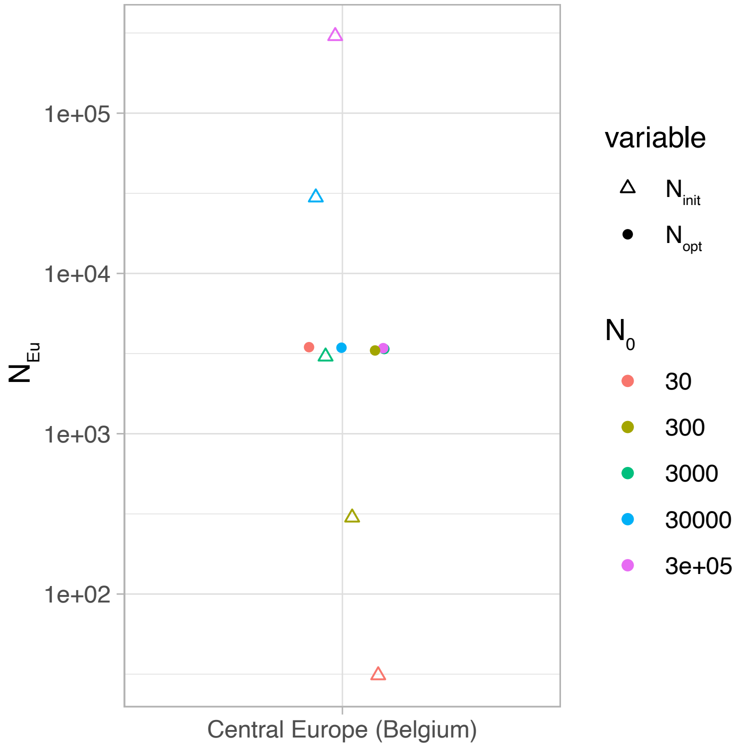


**Figure M8. Convergence of population size of Europe ($N_{Eu}$) regardless of the initial state.** The initial states of population size ($N_0$) were set to 30, 300, 3000, 30000, or 300000, represented in hollow triangles. The optimized $N_{Eu}$ ($N_{opt}$) converged at 3339.5, as represented in solid circles.

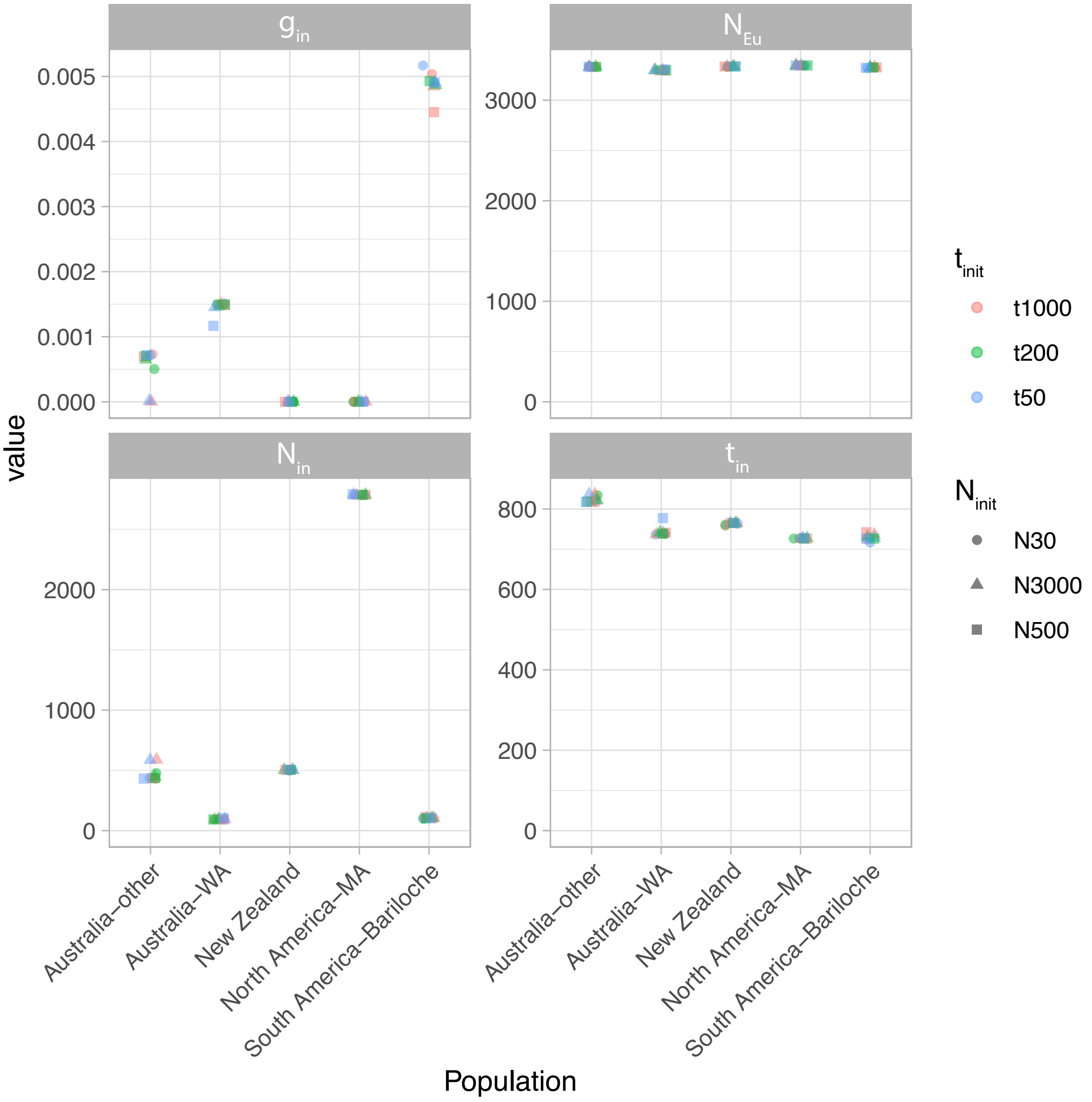


**Figure M9. Convergence of Model A1 regardless of the initial states.** The initial states of population size of introduced population $N_{in}$ and time of divergence from Europe to introduced population $t_{in}$ were explored from 30 to 3000, and 50 to 1000, respectively. The optimized parameters $g_{in}$, $N_{Eu}$, $N_{in}$, and $t_{in}$ converged regardless of the combinations of various initial states.

**Table M2.** Likelihood of the best demographic models when scaling of mutation time and generation time.

| Model | mutation rate | Generation time | Likelihood (log) | NEu | Nin1 | Nin2 | g1 | g2 | m | tin1 | tin2 | tm |
|---|---|---|---|---|---|---|---|---|---|---|---|---|
| A1 South America | 1e-06 | 10 | -5807200.4948 | 3324.73 | 3664.982 | | 0.004 | | | 727.96 | | |
| A1 South America | 1e-07 | 1 | -5807200.4948 | 33247.32 | 36649.82 | | 0.004 | | | 727.96 | | |
| A1 North America | 1e-06 | 10 | -5775459.5465 | 3345.048 | 2786.423 | | 0 | | | 727.13 | | |
| A1 North America | 1e-07 | 1 | -5775459.5465 | 33450.48 | 27864.23 | | 0 | | | 727.13 | | |
| B3 New Zealand and Australia | 1e-06 | 10 | -8408781.1303 | 3257.736 | 456.245 | 975.90 | 0.0 | 0.0032 | 0.373 | 685.15 | 703.53 | 274.59 |
| B3 New Zealand and Australia | 1e-07 | 1 | -8408781.1303 | 32577.36 | 4562.45 | 9759.0 | 0.0 | 0.0032 | 0.373 | 685.15 | 703.53 | 274.59 |
| B2 Australia other and WA | 1e-06 | 10 | -7465490.5904 | 3293.127 | 1718.719 | 2859.6 | 0.002 | 0.0728 | | 747.68 | 89.007 | |
| B2 Australia other and WA | 1e-07 | 1 | -7465490.5904 | 32931.27 | 17187.19 | 28596 | 0.002 | 0.0728 | | 747.68 | 89.007 | |

#### References


**Arita I**. **1979**. The mechanism of spontaneous dedikaryotization in hyphae of Pholiota nameko. *Mycologia* **71**: 603–611.

**Chang CC, Chow CC, Tellier LC, Vattikuti S, Purcell SM, Lee JJ**. **2015**. Second-generation PLINK: rising to the challenge of larger and richer datasets. *Gigascience* **4**: 7.

**Douhan GW, Vincenot L, Gryta H, Selosse M-A**. **2011**. Population genetics of ectomycorrhizal fungi: from current knowledge to emerging directions. *Fungal Biology* **115**: 569–597.

**Fukumasa-Nakai Y, Matsumoto T, Komatsu M**. **1994**. Dedikaryotization of the shiitake mushroom, Lentinula edodes by the protoplast regeneration method. *Journal of General and Applied Microbiology* **40**: 551–562.

**Ginterova A**. **1973**. Dedikaryotization of higher fungi in submerged culture. *Folia Microbiol (Praha)* **18**: 277–80.

**Kamm J, Terhorst J, Durbin R, Song YS**. **2020**. Efficiently inferring the demographic history of many populations with allele count data. *Journal of the American Statistical Association* **115**: 1472–1487.

**Shaw PJA, Kibby C, Mayes J**. **2003**. Effects of thinning treatment on an ectomycorrhizal succession under Scots pine. *Mycological Research* **107**: 317–328.

**Skrede I, Murat C, Hess J, Maurice S, Sonstebo JH, Kohler A, Barry-Etienne D, Eastwood D, Hogberg N, Martin F, *et al.*** **2021**. Contrasting demographic histories revealed in two invasive populations of the dry rot fungus *Serpula lacrymans*. *Molecular Ecology* **30**: 2772–2789.
